# Rab27a co-ordinates actin-dependent transport by controlling organelle-associated motors and track assembly proteins

**DOI:** 10.1101/314153

**Authors:** Noura Alzahofi, Christopher L. Robinson, Tobias Welz, Emma L. Page, Deborah A. Briggs, Amy K. Stainthorp, James Reekes, David A. Elbe, Felix Straub, Edward W. Tate, Philip S. Goff, Elena V. Sviderskaya, Marta Cantero, Lluis Montoliu, Maryse Bailly, Eugen Kerkhoff, Alistair N. Hume

## Abstract

Cell biologists generally consider that microtubules and actin play complementary roles in long- and short-distance transport in animal cells. On the contrary, using melanosomes of melanocytes as a model, we recently discovered that motor myosin-Va, works with dynamic actin tracks, to drive long-range organelle dispersion in microtubule depleted cells. This suggests that in animals, as in yeast and plants, myosin/actin can drive long-range transport. Here we show that SPIRE1/2 and formin-1 (FMN1) proteins generate actin tracks required for myosin-Va-dependent transport in melanocytes. Moreover we show that, in addition to melanophilin/myosin-Va, Rab27a can recruit SPIRE1/2 to melanosomes, thereby integrating motor and track assembly activity at the organelle membrane. Based on this we suggest a model in which organelles and force generators (motors and track assemblers) are linked forming a cell-wide network that allows their collective activity to rapidly disperse the population of organelles long-distance throughout the cytoplasm.

## Introduction

In animal cells, in contrast to plants and yeast, it is generally considered that microtubules (MTs) and actin filaments (AFs) regulate transport in a manner akin to the infrastructure of a developed nation ^1-4^. This ‘highways and local roads’ model suggests that MTs are tracks for long-range transport (highways) between the cell centre and periphery, driven by kinesin and dynein motors. Meanwhile AFs (local roads) and myosin motors work down-stream picking up cargo at the periphery and transporting it for the ‘last μm’ to its final destination. This model makes intuitive sense as MTs in animal cells in culture typically form a polarised radial network of tracks spanning >10 μm from the centrally located centrosome to the periphery and appear ideally distributed for long-distance transport. Meanwhile, with some exceptions in which AFs form uniformly polarised arrays, e.g. lamellipodia, filopodia and dendritic spines, AF architecture appears much more complex. In many fixed cells AF appear to comprise populations of short (1-2 μm length), with random or anti-parallel filament polarity, and not an obvious system of tracks for directed transport ^5,6^.

This view is exemplified by the co-operative capture (CC) model of melanosome transport in melanocytes ^7,8^. Skin melanocytes make pigmented melanosomes and then distribute them, via dendrites, to adjacent keratinocytes, thus providing pigmentation and photo-protection (reviewed in ^9^). The CC model proposes that transport of melanosomes into dendrites occurs by sequential long-distance transport from the cell body into dendrites along MTs (propelled by kinesin/dynein motors), followed by AF/myosin-Va dependent tethering in the dendrites. Consistent with this, in myosin-Va-null cells melanosomes move bi-directionally along MTs into dendrites, but do not accumulate therein, and instead cluster in the cell body ^7,10^. This defect results in partial albinism in mammals due to uneven pigment transfer from melanocytes to keratinocytes (e.g. *dilute* mutant mouse and human Griscelli syndrome (GS) type I patients; Figure 1A) ^11,12^. Subsequent studies revealed similar defects in mutant mice (and human GS types II and III patients) lacking the small GTPase Rab27a (*ashen*) and its effector melanophilin (Mlph) (*leaden*) which recruit and activate myosin-Va on melanosomes ^8,13,14^.

**Figure 1.**
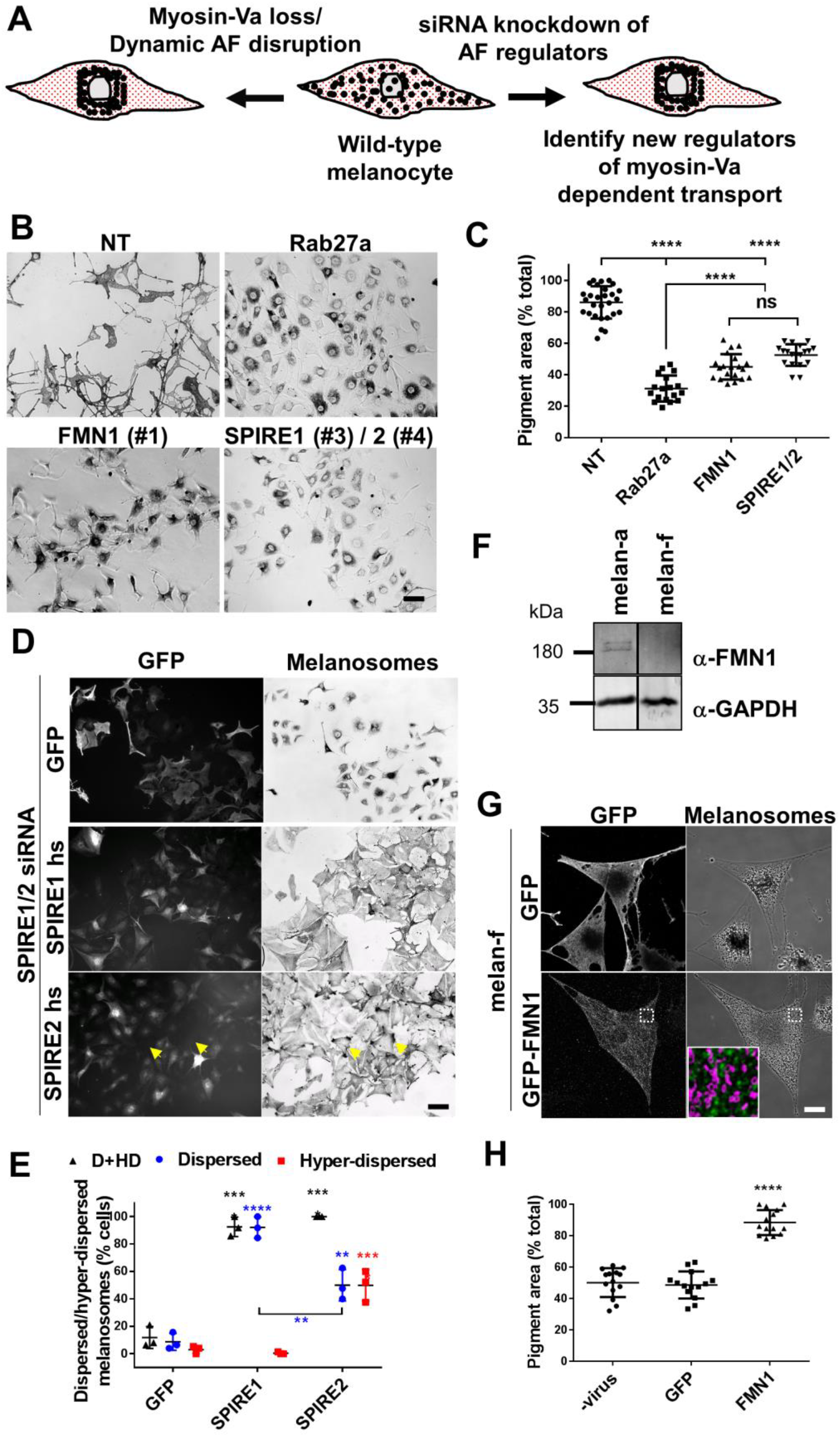
FMN1 and SPIRE1/2 deficient melanocytes show perinuclear melanosome clustering. A) A schematic outlining the strategy used to identify novel regulators of myosin-Va/dynamic AF-dependent melanosome transport. In wild-type melanocytes melanosomes are dispersed throughout the cytoplasm. Genetic loss of myosin-Va (or its regulatory proteins e.g. Rab27a) or pharmacological disruption of a dynamic AFs blocks melanosome dispersion resulting in perinuclear melanosome clustering ^15^. To identify AF associated proteins working with myosin-Va we used siRNA to individually deplete known AF regulators in wild-type melanocytes and screened for those whose depletion pheno-copied the loss of myosin-Va/depletion of dynamic AFs i.e. caused perinuclear clustering of melanosomes. B) melan-a cells were transfected with the indicated siRNA fixed 72 hours cells later and imaged using bright-field optics to observe melanosome distribution (see Experimental Procedures). C) A scatter-plot showing the extent of melanosome dispersion in individual melan-a cells transfected with the indicated siRNA (n = 28 (NT), 18 (Rab27a), 20 (FMN1) and 22 (SPIRE1/2). n.s. indicates no significant difference as determined by one-way ANOVA. All other comparisons yielded highly statistically significant differences p =< 0.0001. D) melan-a cells were transfected with the indicated siRNA. After 72 hours cells were infected with adenovirus expressing GFP or GFP-SPIRE1/2 (human), fixed 24 hours later and processed for immunofluorescence (see Experimental Procedures). Cells were then imaged using bright-field and fluorescence optics to observe melanosome and GFP distribution. E) A scatter-plot showing the percentage of SPIRE1/2 depleted/adenovirus infected melan-a cells in low magnification (10x) fields of view in which melanosomes were dispersed and hyper-dispersed into the peripheral cytoplasm. F) A western blot showing the expression of FMN1 and GAPDH (loading control) in whole cell lysates of melan-a and melan-f melanocytes. G) Confocal micrographs showing the distribution and effect of GFP-FMN1 expression on melanosome distribution in melan-f cells. White dotted boxes in images indicates the region shown in high magnification overlay image (GFP-FMN1 in green and melanosomes in magenta) (see Experimental procedures). H) A scatter plot showing the extent of melanosome dispersion in adenovirus infected melan-f cells expressing the indicated proteins (n = 14 for all conditions). Scale bars represent 50 μm (B and D) and 10 μm (G). (E and H) ****, *** and ** indicates statistical significance of differences p = < 0.0001 p = < 0.001 and p = < 0.01 as determined by one-way ANOVA. Significance indicators above dataset indicate differences compared with GFP control. No other significant differences were observed.

In previous work we tested the CC model by using cell normalisation technology to quantitatively examine the contribution of MTs and AF/myosin-Va to melanosome transport ^15^. Surprisingly, our results indicated that MTs are essential for perinuclear clustering but not peripheral dispersion of melanosomes. Instead we found that MTs retard dispersion, which is dependent upon myosin-Va and a population of dynamic AFs. Functional analysis of mutant proteins suggested that myosin-Va works as a processive motor dispersing melanosomes along AFs whose +/barbed ends are oriented towards the cell periphery. Finally, we developed an activatable myosin-Va, and used this to directly monitor melanosome dispersion in myosin-Va-null cells in real-time. This revealed that myosin-Va can disperse melanosomes rapidly (~1 μm/min) into peripheral dendrites (>10 μm) even in MT depleted cells. Overall our data highlighted the role of myosin-Va and dynamic AFs in long-range transport, rather than tethering, and suggest that melanosome distribution is determined by the balance of MT-dependent clustering and long-range AF/myosin-Va-dependent dispersion. However, studies of AFs organisation in melanocytes have not revealed the existence of a polarised network that would seem requisite for myosin-Va-driven transport and melanosome dispersion ^7^. Thus, the mechanism of this process remained unclear.

Here we investigated this issue and identify SPIRE1/2 and formin-1 (FMN1) AF assembly proteins as essential regulators of myosin-Va driven melanosome dispersion. FMN1 is one of 15 mammalian formins that elongate unbranched AFs ^16,17^. FMN1 and FMN2 comprise the FMN subfamily of formins ^18^. FMN1 function is linked to limb development, while FMN2 acts in oocyte development and memory and learning ^19-22^. Like other formins, FMNs contains two formin homology domains (FH1 and FH2) which drive AF assembly ^17^. Dimeric FH2 forms a ring that encircles the +/barbed ends of AFs, while the proline rich FH1 interacts with G-actin in complex with profilin promoting filament elongation ^23-25^. FMNs are distinct from Diaphanous related formins (Drfs) in that they lack Rho GTPase regulated DID/DAD motifs flanking the active FH1/2 domains ^17,26^. Instead the FMNs are characterised by a short (~30aa) C-terminus FH2 tail which mediates the interaction with SPIRE proteins (termed FSI; <u>F</u>ormin/<u>S</u>PIRE interaction sequence)^27-29^. The N-terminal regions of the FMN subgroup formins are very diverse and no conserved sequence motifs have been identified to date.

SPIRE1/2 are modular proteins that appear to play overlapping roles in the nucleation of AFs at organelle membranes ^18,21,22,30,31^. In line with this SPIRE1/2 contain an N-terminus AF nucleation module, comprising a KIND (kinase non-catalytic C-lobe domain) that interacts with the FSI motif of FMN1/2, and four G-actin binding WH2 (WASP-homology 2) domains ^27,32,33^. This is coupled to a C-terminus membrane binding domain, comprised of SB (SPIREbox, conserved among SPIRE proteins) and FYVE-type zinc finger (Fab1p, YOTB, Vac1, and EEA1) domains ^30,31^. Thus, FMN and SPIRE1/2 proteins may collaborate to assemble AFs at organelle membranes ^21,22^. Previously their combined function has been implicated in regulating oocytes development and repair of DNA damage ^21,34,35^. In mouse oocytes SPIRE1/2 and FMN2 generate AFs for myosin-Vb driven cortical transport of Rab11 vesicles ^21,22^. More recently work has identified a myosin-V globular tail domain binding motif (GTBM) located between the N- and C-terminus modules of SPIRE1/2, which may co-ordinate the recruitment of myosin-V and AF assembly to Rab11 positive intracellular membranes ^36^.

Here we present evidence that the myosin-Va mediated melanosome transport/dispersion in melanocytes is dependent upon AF assembly activities of FMN1 and SPIRE1/2, and that SPIRE1/2 can be recruited to melanosomes by Rab27a. Based on these findings we propose a novel cargo-driven model of organelle dispersion in which Rab27a plays a central role co-ordinating the function of both AF motors and assembly proteins.

## Results

### Formin-1 and SPIRE1/2 proteins are required for melanosome dispersion in melanocytes

To better understand myosin-Va/dynamic AF based dispersive melanosome transport we used siRNA knockdown to test the involvement of known AF regulatory proteins in this process. For this we transfected wild-type melanocytes (melan-a) with an siRNA mini-library comprised of 120 pools (4 target-specific oligonucleotides/pool) directed against the transcripts of known AF regulators. We then visually screened the transfected cells to identify siRNA that induced melanosome clustering, reasoning that knockdown of proteins working with myosin-Va/dynamic AF should result in transport defects like those seen in myosin-Va/dynamic AF deficient cells (Figure 1A). Consistently (5/5 library transfections) we found that knockdown of FMN1 and its interacting partners SPIRE1/2 (double knockdown) induced melanosome clustering similar to that seen for Rab27a knockdown (part of a myosin-Va receptor at the melanosome), albeit that the extent of clustering was significantly lower (Figure 1B-C; mean pigment area (% total); NT = 86 +/- 10.28 %, Rab27a = 31.28 +/- 8.242 %, FMN1 = 45.07 +/- 8.061 % and SPIRE1/2 = 52.56 +/- 6.868 %). The efficacy of mRNA knockdown was confirmed using quantitative real-time-PCR (Q-RT-PCR) (Figure S1A-C). In addition we used RT-PCR to compare the levels of expression of SPIRE1/2 and FMN1/2 mRNA in melan-a cells (Figure S1D). Consistent the lack of effect of siRNA transfection, by this approach we could not detect FMN2. Meanwhile quantification showed that SPIRE1 mRNA expression exceeded that of SPIRE2 by a factor of five and that FMN1 expression was not comparable to the sum of SPIRE1/2 expression (Figure S1D; mRNA copies (x103)/50ng total RNA; SPIRE1 = 17.33 +/- 1.883, spire2 = 3.468 +/- 0.5726, SPIRE1/2 20.8 +/- 1.931 and FMN1 = 27.2 +/- 4.17).

Interestingly, although single transfection with SPIRE1 and SPIRE2 specific siRNA was effective in reducing mRNA only SPIRE1 knockdown resulted in a significant decrease in the proportion of cells with dispersed melanosomes compared with control (NT) siRNA transfected cells (Figure S1B-E; dispersed melanosomes (% cells); NT = 86.43 +/- 4.887 % versus SPIRE1 = 46.93 +/- 10.64 % and SPIRE2 = 82.57 +/- 13.26 %). Nevertheless, the effect of SPIRE2 depletion could be seen by the significantly lower proportion of cells with dispersed melanosomes seen in SPIRE1/2 versus SPIRE1 alone siRNA transfections (Figure S1B-F; mean % of cells with dispersed melanosomes; SPIRE1/2 = 12.73 +/- 5.878 %). Also, in a small, but significant, subset of SPIRE1 depleted cells we observed that melanosomes accumulated in the cortical cytoplasm, hereafter termed ‘hyper-dispersed’, that we did not observe in SPIRE2 or SPIRE1/2 depleted cells (Figure S1E-F; mean hyper-dispersed melanosomes (% total cells) = 6.324 +/- 2.645 %). This is consistent with the higher expression of SPIRE1 versus SPIRE2 mRNA and indicates that there may be subtle differences in the function of the SPIRE1/2 proteins in melanocytes (Figure S1D).

To confirm the results of FMN1 knockdown we generated immortal FMN1 deficient melanocytes (melan-f) from FMN1^pro^ mutant mice (see Experimental procedures) ^37^. Consistent with the results of knockdown melan-f cells manifested perinuclear melanosome clustering (Figure 1G-H). Interestingly, the extent of melanosome clustering in melan-f and FMN1 depleted melan-a was similar to one another and to that seen in melan-a cells depleted of dynamic AFs using the G-actin sequestering drug latrunculin A (Figure S2, 1G; mean pigment area (% total); melan-a with 30 nM latrunculin A = 51.56 +/- 12.14 %, versus melan-f cells with 0 nM latrunculin A = 45.48 +/- 11.9 %)^15^. Moreover, melanosome distribution in melan-f cells was not significantly affected by incubation with 30 nM latrunculin A, previously found to deplete the dynamic AF pool required for melanosome dispersion (mean pigment area (% total); melan-f with 30 nM latrunculin A = 45.89 +/- 13.29 %). This indicates that FMN1 (and SPIRE1/2) assemble dynamic AFs that, with myosin-Va, are essential for melanosome dispersion.

To test the specificity of melanosome clustering in SPIRE1/2 and FMN1 deficient cells we generated adenoviruses that express human (hs) SPIRE1/2 (these are resistant to mouse-specific siRNA) and mouse FMN1 as fusions to the C-terminus of EGFP. Add-back experiments in SPIRE1/2 depleted melan-a and melan-f melanocytes using these vectors confirmed that melanosome clustering was due to depletion of the endogenous proteins (Figure 1D-H; E, mean % of cells with dispersed and hyper-dispersed (D+HD) melanosomes; SPIRE1/2 knockdown - virus = 11.03 +/- 6.668 %, GFP = 11.83 +/- 8.02 %, SPIRE1 = 92.58 +/- 7.072 %, SPIRE2 = 94.67 +/- 11.36 %; H, mean pigment area (% total), – virus = 50.15 +/- 9.228 %, GFP = 48.66 +/- 8.618 %, FMN1 = 88.43 +/- 7.972 %). Interestingly, among significant subset of cells rescued by SPIRE2, but not SPIRE1, we observed hyper-dispersion of melanosomes, similar to that seen in cells singly depleted of SPIRE1 (Figure 1D, S1D-F (arrows in images); mean % of cells with hyper-dispersed melanosomes; GFP = 3.088 +/- 2.748 %, SPIRE1 = 0.4695 +/- 0.8132 %, SPIRE2 = 49.96 +/- 11.44 %). This effect was particularly apparent in cells expressing relatively low levels of protein. Altogether these results indicate that FMN1 and SPIRE1/2 function with myosin-Va to disperse melanosomes in melanocytes and further underline the imperfect functional overlap of SPIRE2 and SPIRE1.

### The membrane binding domain of SPIRE1/2 is related to the Rab binding domain of Rab3/27 effectors

Previous studies reported that the sequence of the conserved SB portion of SPIRE1 is similar to the Rab binding domain (RBD) of the Rab3/27 effector Rabphilin-3A, and more recently the C-terminus membrane binding (SB and FYVE) portion of SPIRE1 was found to interact with Rab3a *in vitro* ^30,38^. The N-terminus RBD of Rabphilin-3A is composed of two α-helical domains (termed SHD (<u>s</u>ynaptotagmin <u>h</u>omology <u>d</u>omain) 1 and 2) that flank a FYVE-type zinc finger motif ^39^. Similar structures are found in other Rab27 effectors, including Mlph which regulates melanosome dispersion via activation of myosin-Va ^8,13,40,41^. The N-terminus SHD1 and 2 of Mlph make direct and essential contacts with Rab27 ^40-42^. An alignment of vertebrate SPIRE1/2 sequences encoding the SB, the FYVE-type domain and C-terminal flanking sequences with the RBDs of vertebrate Mlph and the related Rab27 effector Myrip (<u>my</u>osin/<u>R</u>ab interacting <u>p</u>rotein) revealed that these are closely related (Figure S3A). In particular, residues which make contact with Rab27 are conserved between Mlph and SPIRE1/2 suggesting that these proteins interact with Rab27 via similar mechanisms ^41^. Phylogenetic analysis of the sequences of the putative RBDs of SPIRE proteins with those of other SHD containing Rab3/27 effectors further confirmed the close relationship between these groups. This analysis also grouped SPIRE1/2 into the family of SHD containing Rab3/27 effectors (Figure S3B).

### SPIRE1/2 interact with GTP-bound, active Rab27a

The sequence similarities of SPIRE1/2 to SHD-family proteins suggest that SPIRE1/2 could be Rab27 effectors nucleating AFs at the melanosome membrane in melanocytes. As a first step to test this we performed GST pull-down experiments using bacterially expressed, purified GST-Rab27a (GTP-locked Rab27a-Q78L and GDP-locked Rab27a-T23N mutant proteins) and lysates of HEK293 cells transiently expressing Myc-epitope tagged-SPIRE1/2 (Figure 2B). By Western blotting we found that GST-Rab27a-Q78L pulled-down greater quantities of Myc-tagged SPIRE1/2 compared with GST-Rab27a-T23N, supporting the possibility that SPIRE1/2 are novel Rab27 effector proteins (Figure 2B-C). Also, we observed that more SPIRE1 than SPIRE2 was pulled-down, indicating that SPIRE1 has a higher affinity for Rab27a than SPIRE2 (Figure 2B-C). This is consistent with the higher level of sequence similarity between Mlph/Myrip and SPIRE1 versus SPIRE2 (Figure S3A).

**Figure 2.**
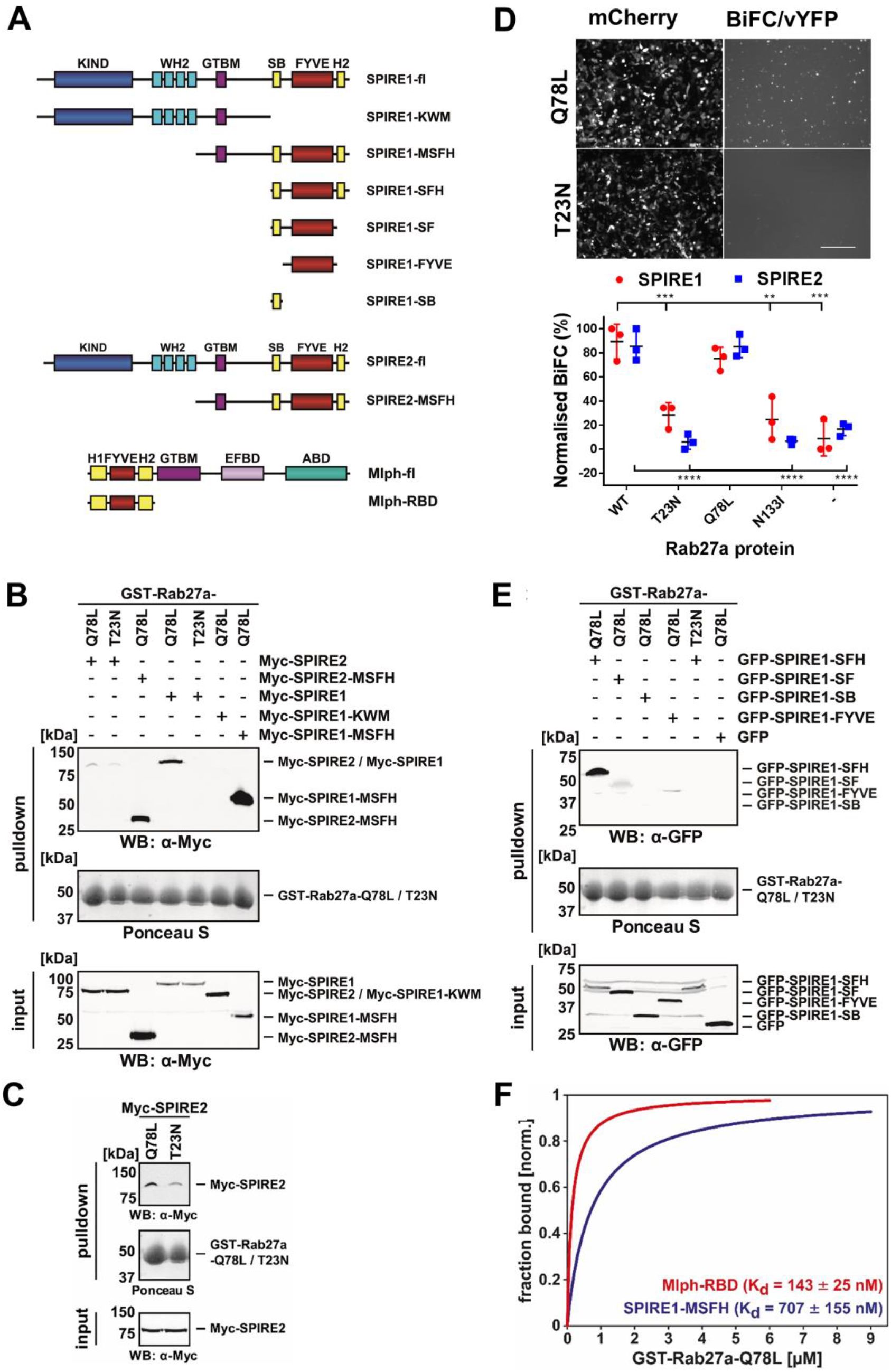
SPIRE1/2 interact with active Rab27a via their membrane binding C-termini. A) A schematic representation of the domain structure of SPIRE1/2 and Mlph, and the correspondence with truncations used in interaction studies (B-E). Interaction of SPIRE1/2 and Rab27a was investigated using GST-pull-down (B, C, E and F) and BiFC (D) assays (as described in Experimental procedures). B, C and E) Western blots and Ponceau S stained filters showing the results of pull-down assays measuring the interaction of GST-Rab27a (active Q78L and inactive T23N mutants) with SPIRE1/2 and the indicated truncations (Myc-tagged (B, C) and GFP-tagged (E)) *in vitro*. C) is a contrast enhanced version of the section of B) showing interaction of SPIRE2 with Rab27a-Q78L and Rab27a-T23N. D) Fluorescence images and a scatter plot (upper and lower panels) showing the results of the BiFC assay reporting the interaction of Rab27a and SPIRE1 in HEK293a cells (see Experimental procedures). Images of mCherry indicate transfection efficiency and vYFP indicates BiFC i.e. interaction. The scatter plot shows the BiFC signal for populations of cells expressing SPIRE1/2 with active and inactive Rab27a mutants. Data shown are from 3 independent experiments. ****, *** and ** indicate significant differences of p = < 0.0001, p = < 0.001 and p = <0.01 as determined by one-way ANOVA of data for SPIRE1/2 with different Rab27a proteins. No other significant differences were observed. Two-way ANOVA comparison of BiFC signal for the Rab27a proteins with different SPIREs revealed no significant differences. F) Line plots showing the extent of binding of GFP-SPIRE1-MSFH (n = 4) and GFP-Mlph-RBD (n = 3) as a function of increasing GST-Rab27a-Q78L concentrations. Error bars represent SEM and the equilibrium dissociation constants (*K_d_*) are provided. *K*, KIND; *W*, WH2; *M*, GTBM, globular tail domain binding motif; *S*, SB, SPIRE-box; *F*, FYVE-type zinc finger; *H*, C-terminal flanking sequences similar to H2 of SHD-proteins; *WB*, Western blotting.

To further investigate Rab27a:SPIRE1/2 interaction we used a bimolecular fluorescence complementation (BiFC) assay in which Rab27a and SPIRE1/2 were transiently co-expressed in HEK293 cells as fusions to C-terminus and N-terminus halves of the Venus yellow fluorescent protein (vYFP). We observed significantly higher BiFC signal in cells transfected with wild-type Rab27a and the Q78L mutant compared with the inactive mutants T23N and N133I in which the mean BiFC signal was similar to cells transfected with the vYFP fragments alone (Figure 2D; mean normalised BiFC (% max); SPIRE1/SPIRE2 WT = 89.35/85.41 %, T23N = 28.35/5.969 %, Q78L = 75.21/85.23 %, N133I = 24.71/6.601 % and vYC alone = 8.637/16.65 %). These data concur with the results of pull-down assays and further indicate that SPIRE1/2 are novel Rab27a effectors.

### The C-terminus membrane binding module of SPIRE1/2 interacts with Rab27a

To map the Rab27a binding site(s) within SPIRE1/2 we tested the interaction of SPIRE1 deletion mutants using the GST-pull-down assay. As indicated above SPIRE1/2 proteins have the following domain organisation: KIND (K), WH2-cluster (W); GTBM (M), SPIRE-box (S), FYVE-type (F), C-terminal flanking sequences similar to helix 2 of SHD-proteins (H) (Figure 2A). Consistent with the distribution of sequence similarity between SPIRE1/2 and other Rab27 effectors, we found a strong interaction of GST-Rab27a-Q78L with Myc-SPIRE1-MSFH (and Myc-SPIRE2-MSFH) proteins but not Myc-SPIRE1-KWM (Figure 2B). To further characterise the Rab27a binding site in SPIRE1 we generated a series of truncation mutants and tested their interaction with Rab27a-Q78L. We found that SPIRE1-SFH interacted strongly with Rab27a-Q78L and that the FYVE-only and SF proteins also interacted, albeit to a lesser extent (Figure 2A, E). These observations indicate that the C-terminus SFH fragment, which contains elements conserved with the RBD of other Rab27a effectors, interacts with Rab27a and suggests that SPIRE1/2 interact with Rab27a via a mechanism similar to other effectors e.g. Mlph.

### SPIRE1/2 interact with Rab27a with lower affinity than Mlph

Rab27 effectors have been classified into high (e.g. rabphilin, Noc2, Slp2-a) and low affinity groups (e.g. Mlph, Myrip). In line with this the dissociation constant (K_d_) for the Rab27a-Q78L interaction with the Slp2-a or Mlph RBDs were determined to be 13.4 and 112 nM respectively ^43^. It has been proposed that the distinct Rab27a binding activities ensure the order in which the effectors function in transport processes ^43^. To compare the interaction of Rab27a with SPIRE1/2 to that of other effectors e.g. Mlph, we quantified the affinity of interaction of GFP-SPIRE1/2-MSFH and GFP-Mlph-RBD with GST-Rab27a-Q78L in pull-down assays by measurements of the fluorescent protein depletion from HEK293 cell lysates (Figure 2A, F). This revealed dissociation constants of 143 (+/- 25) nM and 707 (+/- 155) nM and maximum binding levels of 39.4% and 12.0% for the GFP-Mlph-RBD and GFP-SPIRE1-MSFH with GST-Rab27a-Q78L. These data, together with previous estimates of the affinity of Mlph:Rab27a interaction (see above) indicate that SPIRE1 is a weaker Rab27a interactor than Mlph ^43^. Using this approach we were unable to quantify the SPIRE2:Rab27a interaction due to the very low affinity of SPIRE2 for Rab27a-Q78L. This further suggests that Rab27a:SPIRE1 interaction is stronger than Rab27a:SPIRE2 interaction (Figure 2B, C, F).

### SPIRE1/2 associate with melanosomes by a Rab27a-dependent mechanism

The above findings indicate that SPIRE1/2 are Rab27 effectors whose expression is required for dispersion of melanosomes into the peripheral dendrites of melanocytes. Given that Rab27a is present on the cytoplasmic face of the melanosome membrane, this suggests that SPIRE1/2 may associate with melanosomes in a Rab27a-dependent manner. To test this, we expressed GFP-tagged SPIRE1/2 proteins and Rab27a in melan-a cells and used confocal microscopy to examine their intracellular localisation. We observed that all three proteins were distributed in a punctate pattern throughout the cytoplasm (Figure 3A-C). High magnification imaging and intensity profile plots through melanosome containing areas revealed that spots of SPIRE1 and Rab27a often overlapped with melanosomes while this was less obvious for SPIRE2 (Figure 3A-C; Pearson linear correlation coefficient = 0.823 for Rab27a, 0.623 for SPIRE1 and −0.081 for SPIRE2). We also observed that Myc-SPIRE1-MSFH overlapped with melanosome resident protein tyrosinase-related protein 1 (TRP1) and other melanosome targeted proteins (co-expressed mRuby3-Rab27a and GFP-myosin-Va-CC-GTD (that includes the melanocyte-specific coiled coil and globular tail domains; Figure S4)). These observations indicate that SPIRE1 associates with melanosomes.

**Figure 3.**
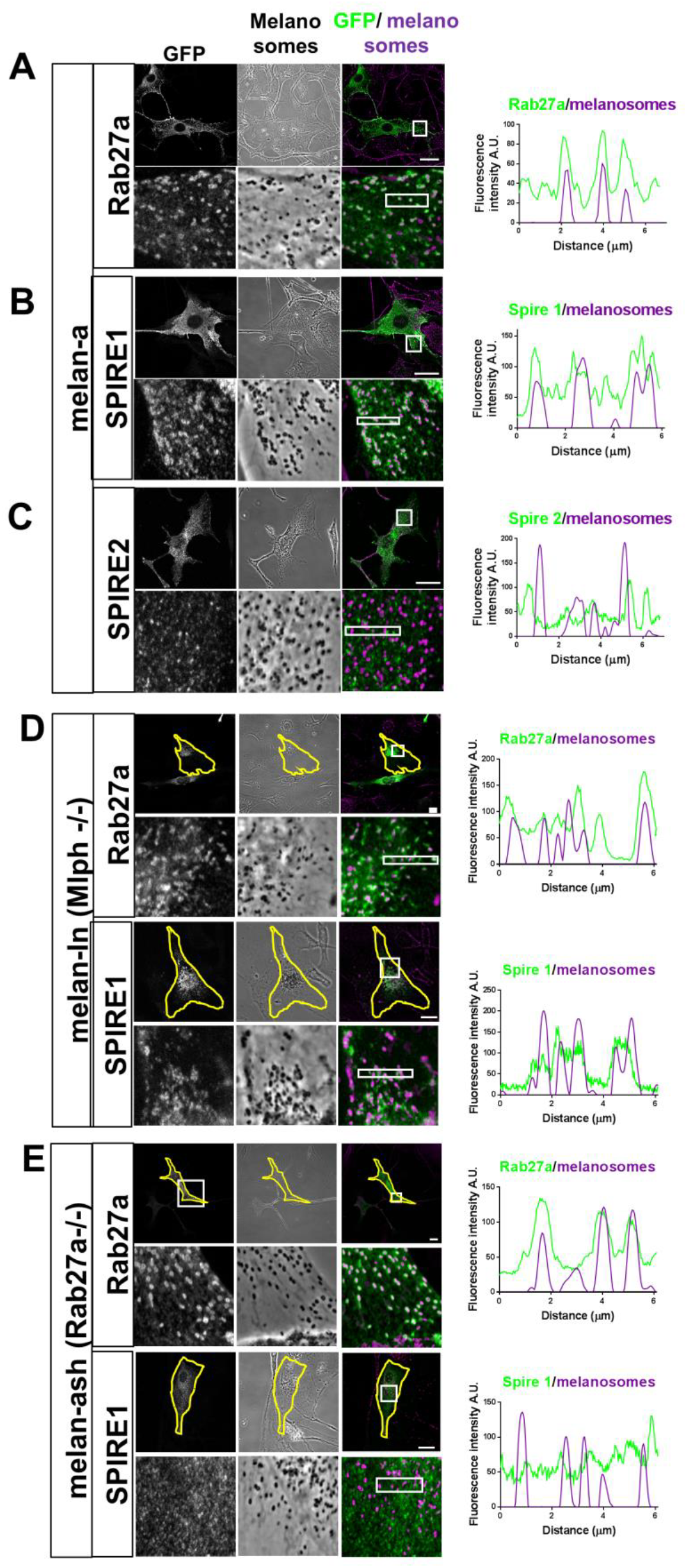
Rab27a recruits SPIRE1 to melanosomes in melanocytes. Melanocytes were transfected with plasmids allowing expression of the indicated proteins as fusions to the C-terminus of EGFP. Cells were fixed after 48 hours, stained with GFP-specific antibodies to detect the expressed proteins and the intracellular distribution of expressed protein and melanosomes was observed using a confocal microscope (see Experimental procedures). A-E) Single confocal z-sections of the distribution of each protein, pigmented melanosomes (transmitted light/phase contract images) and merge images (melanosomes pseudo-coloured magenta). Upper panels show whole cells. Boxes indicate regions shown in lower panels at high magnification allowing comparison of the distribution of melanosomes and fluorescent protein. Line plots are fluorescence intensity profile plots of the boxed regions in high magnification images averaged along the vertical axis. (A-C), (D) and (E) are melan-a, melan-ln and melan-ash cells. D and E) Yellow lines indicate the borders of transfected cells. Scale bars represent 20 μm.

To test whether this was Rab27a-dependent we repeated the above experiment using Mlph- and Rab27a-deficient melanocytes (melan-ln (in which Rab27a is retained on the melanosomes membrane) and melan-ash) ^44,45^. This revealed that while Rab27a associated with melanosomes in both cell types (and as expected dispersed melanosomes in melan-ash cells), SPIRE1 associated with perinuclear clustered melanosomes in melan-ln cells only (Figure 3D-E; Pearson linear correlation coefficient = 0.601 and 0.739 for Rab27a, and = −0.230 and 0.617 for SPIRE1 in melan-ash and melan-ln). These data indicate that association of SPIRE1 with melanosomes is dependent upon Rab27a and supports the hypothesis that SPIRE1 is a Rab27 effector.

Given the sequence similarity between SPIRE1 and SPIRE2 plus in vitro and BiFC interaction data, we were surprised that we did not clearly detect GFP-SPIRE2 at the melanosome membrane (Figure S3A, 2B-D, 3C). Based on the results of GST pull-down experiments we suggest that the apparent weakness of the Rab27a:SPIRE2 versus Rab27a:SPIRE1 interaction might explain this observation (Figure 2B-C). To further investigate Rab27a:SPIRE2 interaction in melanocytes we modified the previously described ‘minimyosin’ protein (that couples an active motor-lever arm (S1) fragment of myosin-Va to the RBD of the Rab27 effector synaptotagmin-like protein 2-a (Slp2-a)) by replacing the Slp2-a RBD fragment with SPIRE1/2 proteins to create ‘myoSPIRE1/2’ fusion proteins (Figure 4A). We then tested the ability of these proteins to disperse clustered melanosomes in melan-ln and melan-ash cells (i.e. in the presence versus absence of melanosome associated Rab27a). In melan-ln we found that both myoSPIREs, like minimyosin but not Rab27a, significantly dispersed melanosomes compared with GFP alone (Figure 4B-C; mean pigment area (% total); GFP = 26.3 +/- 7.43 %, myoSPIRE1 = 63.4 +/- 7.96 %, myoSPIRE2 = 66.6 +/- 3.76 %, mini-Va = 62.3 +/-10.8 % for and Rab27a = 31.3 +/- 10.7 %). In contrast, in melan-ash, although Rab27a expression dispersed melanosomes, neither minimyosin nor myoSPIRE proteins did so to a significantly greater extent than GFP alone (Figure 4C; mean pigment area (% total); GFP = 31.9 +/- 6.80 %, myoSPIRE1 = 37.8 +/- 5.84 %, myoSPIRE2 = 38.6 +/- 5.97 %, mini-Va = 39.2 +/-1.97 % and Rab27a = 66.9 +/- 6.20 %). This supports the hypothesis that both SPIRE1 and SPIRE2 can interact with melanosome associated Rab27a in melanocytes. Consistent with this using confocal microscopy we observed that spots of both myoSPIREs were often located adjacent to melanosomes in melan-ln cells (Figure 4B).

**Figure 4.**
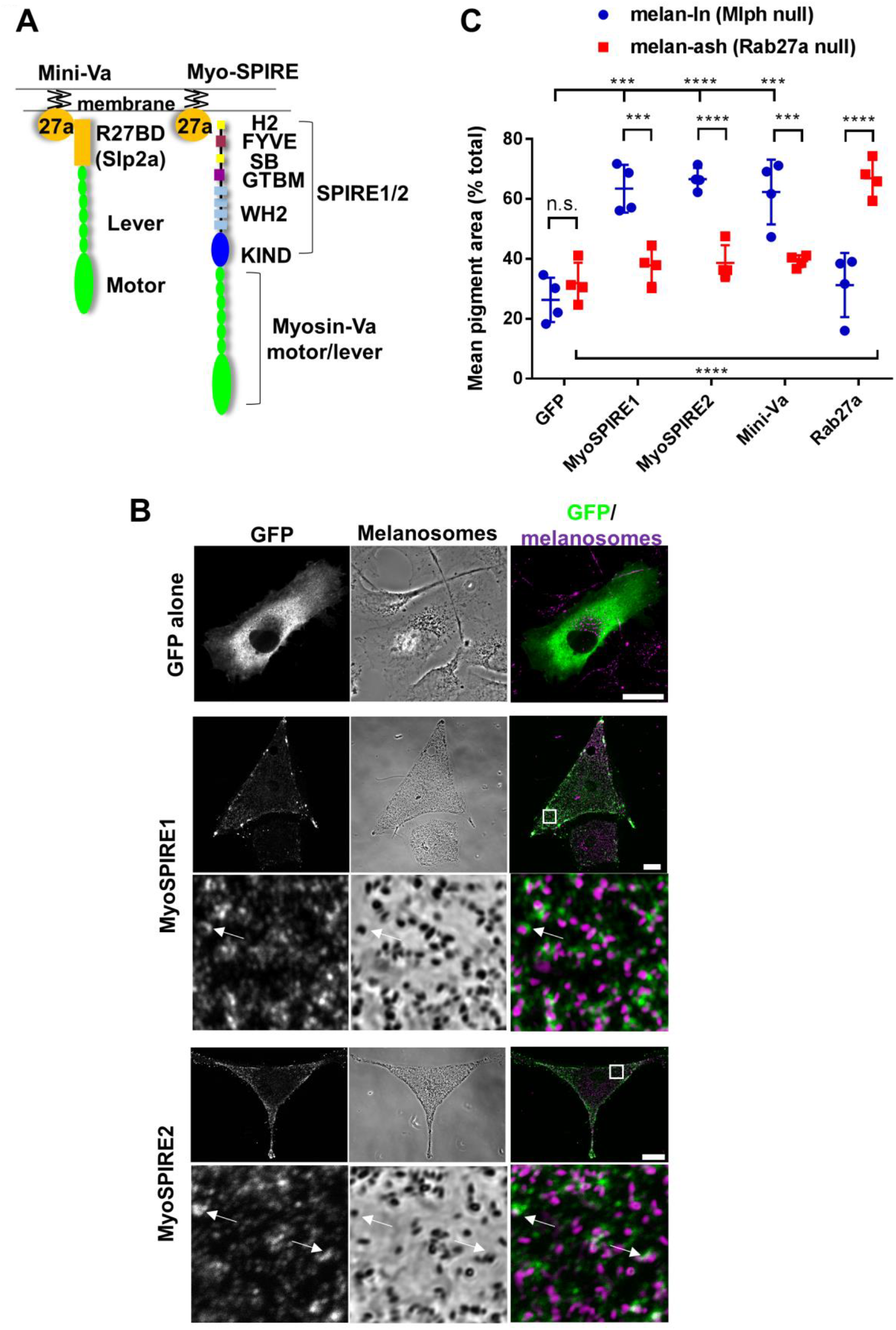
Functional evidence that SPIRE1/2 interact with melanosome associated Rab27a in melanocytes. melan-ln (Mlph null) and melan-ash (Rab27a null) melanocytes were infected with viruses expressing the indicated proteins. Cells were fixed after 24 hours, stained with GFP-specific antibodies and the intracellular distribution of GFP-fusion proteins and melanosomes was observed using a confocal microscope (see Experimental procedures). A) A schematic representation of the structure of mini-Va and myoSPIRE1/2 proteins and their interaction with membrane associated Rab27a. *R27BD*, Rab27 binding domain; *SB*, SPIRE-box; *GTBM*, globular tail binding domain. B) Single confocal z-sections showing the distribution of expressed proteins, melanosomes and their colocalisation (from left to right). In merge images melanosomes are false-coloured magenta. For myoSPIRE1/2 upper and lower panels are low and high magnification images. Boxes in low magnification images indicate the area presented in the high magnification images below. Scale bars represent 10 μm. Arrows in high magnification images indicate examples of co-localisation of spots of myoSPIRE with melanosomes. C) Scatter-plot showing the effect of expression of myoSPIRE and other proteins on melanosome distribution in melanocytes. Data presented are mean pigment area measurements from 4 independent experiments. In each case pigment area was measured for 10 cells expressing each of the different proteins. **** and *** indicate significant difference p =< 0.0001 and p =< 0.001 as determined by one-way ANOVA. n.s. indicates no significant difference.

### The N-terminus AF nucleation module (KIND and WH2) is essential for SPIRE function in melanosome transport

To understand better to role of SPIREs in melanosome transport we investigated the domains of SPIRE1/2 make significant functional contributions. To do this we generated adenoviruses that express the N-terminus KW (FMN interaction and AF nucleation module) and C-terminus MSFH (myosin-V binding and membrane interaction modules) fragments of siRNA resistant human SPIRE2 and tested their ability to rescue melanosome dispersion in SPIRE1/2 depleted melan-a cells (Figure 5A). We found that while neither truncation dispersed melanosomes as efficiently as intact SPIRE2, SPIRE2-KW dispersed melanosomes to a significantly greater extent than SPIRE2-MSFH or GFP-alone (Figure 5B-C; mean % of cells with dispersed and hyper-dispersed (D+HD) melanosomes, KW = 85.8 +/- 3.539 % and MSFH = 10.78 +/- 9.648 %). This suggests that both N- and C-terminus modules are required for optimal SPIRE1/2 function, but that the N-terminus FMN1 interaction and AF nucleation activities are the most significant functional elements of SPIRE1/2 in melanosome transport while the membrane targeting fulfils a secondary role increasing their efficiency by targeting AF nucleation to Rab27a positive melanosomes.

**Figure 5.**
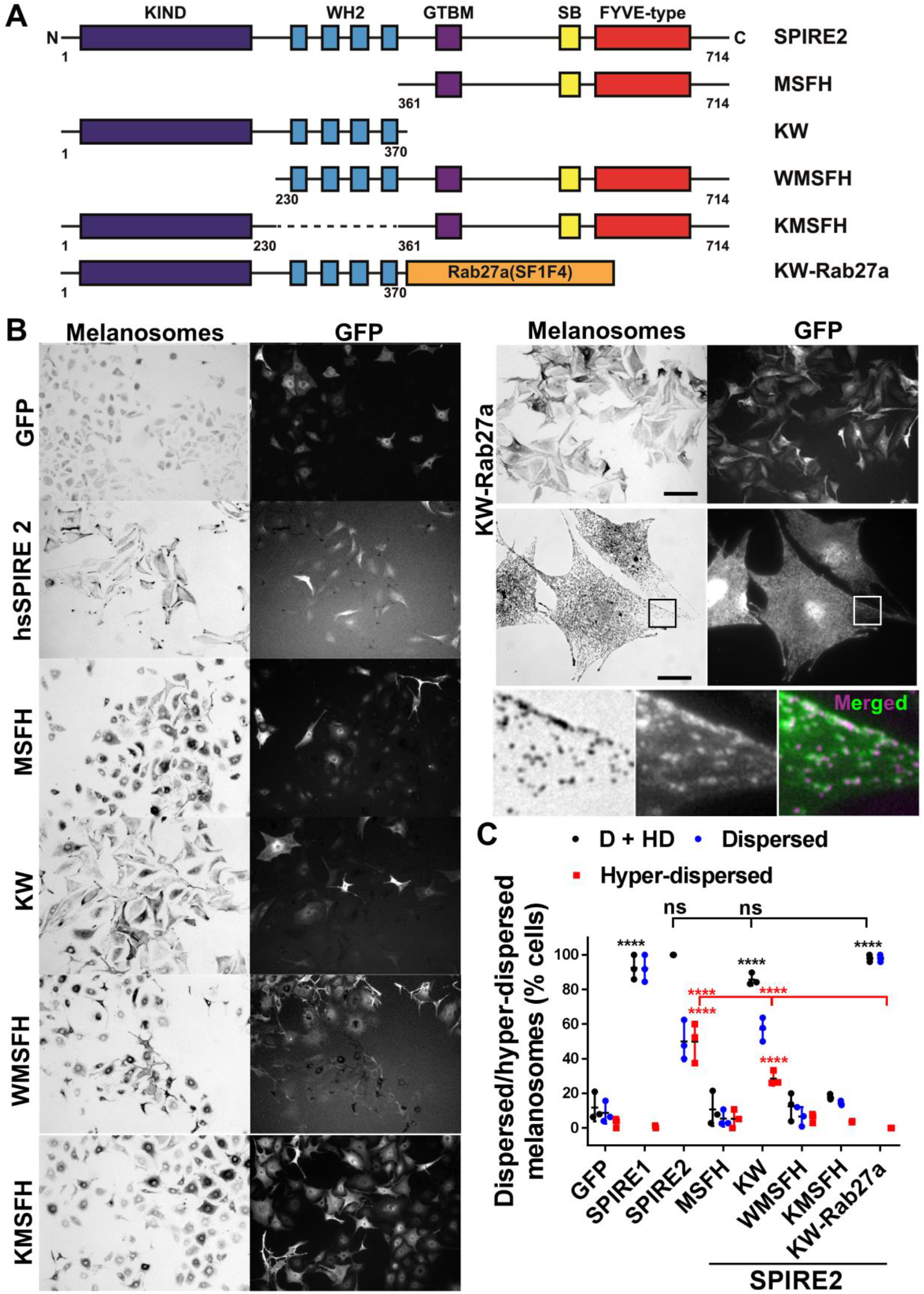
The FMN interaction (KIND) and AF nucleation (WH2) activities of SPIRE2 are essential for melanosomes dispersion. A) A schematic representation of the domain structure of human SPIRE2 and the correspondence with truncations and chimeric proteins used in functional studies (B-C). Numbers indicate amino acid boundaries. *K*, KIND; *W*, WH2; *M*, GTBM, globular tail domain binding motif; *S*, SB, SPIRE-box; *F*, FYVE-type zinc finger; *H*, C-terminal flanking sequences similar to H2 of SHD-proteins. B) melan-a cells were depleted of SPIRE1/2 by siRNA transfection and 72 hours later infected with adenoviruses expressing the indicated proteins. Cells were fixed 24 hours later, processed for immunofluorescence and imaged using bright-field and fluorescence optics to observe melanosome and protein distribution/expression (see Experimental procedures). Scale bars represent 100 μm and 20 μm in low and high magnification images. Boxes in KW-Rab27a images indicate the region shown below. For the merged image green = KW-Rab27a in and magenta = melanosomes. C) Is a scatter plot showing the percentage of human SPIRE2 expressing SPIRE1/2 siRNA depleted melan-a cells in which melanosomes are classed as dispersed and/or hyper-dispersed. **** indicates significant difference p = < 0.0001 as determined by one-way ANOVA. Asterisks indicate the significance of difference of the dataset compared with GFP expressing control unless otherwise indicated. Asterisk colour indicates the class of data tested. No significant differences were seen between other datasets and GFP or KW-Rab27a. Results shown are representative of 4 independent experiments.

To investigate this further we tested WMSFH and KMSFH truncations, which individually lack either the KIND or WH2 domains, respectively. We found that neither protein dispersed melanosomes in SPIRE1/2 depleted cells, indicating that both the AF nucleation (WH2) and FMN interaction (KIND) are essential for SPIRE2 function (Figure 5B-C; mean % of cells with dispersed and hyper-dispersed (D+HD) melanosomes; WMSFH = 12.57 +/- 8.109 % and KMSFH = 17.9 +/- 1.583 %). Similar results were seen in studies in which human SPIRE1 was used to rescue SPIRE1/2 double depletion (Figure S5). Taken with the results of SPIRE1/2:Rab27a interaction studies, the above data indicate that SPIRE1/2 proteins may be melanosome associated AF nucleators that work with FMN1, and are essential for myosin-Va driven melanosome dispersion.

In these experiments we also noted that, in contrast to intact SPIRE1 and SPIRE2, which differ in their ability to promote melanosome hyper-dispersion, KW truncation of both proteins behave similarly (Figure 5C, S5B; mean % of cells with hyper-dispersed melanosomes, SPIRE2-KW = 28.58 +/- 4.119 % and SPIRE1-KW = 36.69 +/- 7.746 %). These observations suggest that difference(s) in the C-terminus membrane binding element of SPIRE1/2 are the basis of their differing ability to disperse melanosomes. Related to this one possibility is that it is the relatively low affinity of SPIRE2:Rab27a versus SPIRE1:Rab27a interaction that allows it to promote hyper-dispersion of melanosomes.

### The FH1, FH2 and FSI domains are required for FMN1 function in melanosome transport

We then investigated FMN1 function in melanosome transport. Using confocal microscopy, we examined the intracellular distribution of GFP-FMN1 in melan-f cells. This revealed that FMN1 was distributed throughout the cytoplasm and was not strongly associated with melanosomes (Figures 1G (inset); 6B). To probe its role in transport we tested the ability of FMN1 mutants to disperse clustered melanosomes in melan-f cells (Figure 6A). We found that expression of a C-terminus fragment, encompassing the conserved FH1 and FH2 domains and the FSI motif (FH1-FH2-FSI), restored peripheral melanosome distribution to the same extent as intact FMN1, while the reciprocal N-terminus fragment did not (Figure 6; mean pigment area (% total); GFP = 50.69 +/- 14.29 %, FMN1 = 89.92 +/- 6.598 %, N-term = 51.44 +/- 10.17 %, FH1-FH2-FSI = 89.28 +/-8.512 %). Truncations that removed the FSI or FH1 were unable to disperse melanosomes in melan-f cells to the same extent as wild-type protein (Figure 6; mean pigment area (% total); ΔFSI = 79.52 +/- 12.34%, FH2-FSI = 42.77 +/- 13.89 %). Also we found that point mutations predicted, based on sequence similarity, to disrupt FH2 contact with AF +/barbed ends (I1074A and K1229D) or the electrostatic interaction of the FSI with SPIRE-KIND (K1418E) significantly reduced FMN1 function in transport (Figure 6; mean pigment area (% total), I1074A = 57.77 +/- 12.73%, K1229D = 77.75 +/- 12.53 and K1418E = 72.1 +/-17.34 %) ^23,29^. These results indicate that the FH1 and FH2 domains and the FSI motif are essential for FMN1 function in transport and indicate that FMN1 works with SPIRE1/2 to assemble AFs in melanocytes but may not stably associate with melanosomes.

**Figure 6.**
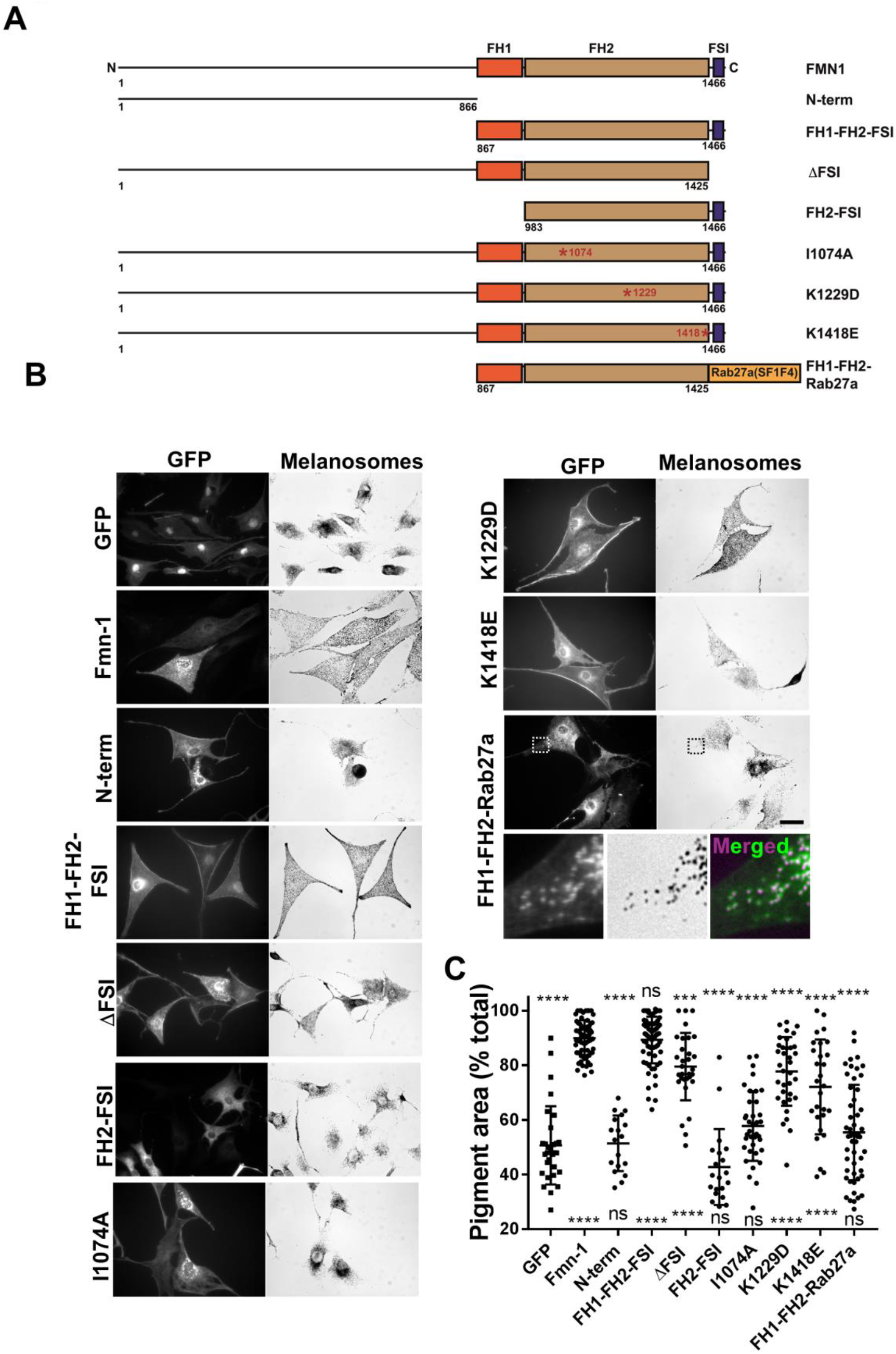
The AF assembly (FH1-FH2) and SPIRE interaction (FSI) domains of FMN1 are essential for melanosome dispersion. A) A schematic representation of the domain structure of murine FMN1 and the composition of truncations and chimeric proteins used in functional studies (B-C). Red asterisks indicates the site of point mutations. Numbers indicate amino acid boundaries. B) melan-f cells were plated on glass cover-slips and infected with adenoviruses expressing the indicated proteins. Cells were fixed 24 hours later, processed for immuno-fluorescence and the intracellular distribution of expressed protein and melanosomes (bright-field) was observed using a fluorescence microscope (see Experimental procedures). Scale bars represent 20 μm. Dashed boxes in FH1-FH2-Rab27a images indicate the region of the image shown in high magnification in the panels below. The merged image show melanosomes false coloured *magenta* and GFP-FH1-FH2-Rab27a in *green*. C) A scatter-plot showing the extent of pigment dispersion in cells expressing the indicated proteins. n = 30 (GFP), 59 (FMN1), 18 (N-term), 70 (FH1-FH2-FSI), 31 (ΔFSI), 20 (FH2-FSI), 37 (I1074A), 35 (K1229D), 28 (K1418E) and 51 (FH1-FH2-Rab27a). ****, ** and * indicate significant differences of p=< 0.0001, p =< 0.01 and p =< 0.05 as determined by one-way ANOVA. n.s. indicates no significant difference. Significance indicators above and below the data indicate differences between the data and the positive (FMN1 wild-type) and negative (GFP alone) controls. Results shown are representative of 3 independent experiments. *FH*, formin homology; *FSI*, Formin-SPIRE interaction sequence.

### A model for myosin-Va/AF dependent melanosome dispersion

The data presented above indicate firstly, that FMN1 and SPIRE1/2 regulate pigment transport in melanocytes, and secondly, that along with myosin-Va, Rab27a can recruit SPIRE1/2 to melanosomes. The latter point raises the interesting possibility that Rab27a drives melanosome dispersion by co-ordinating the activity of both AF dependent motors (i.e. myosin-Va) and AF assembly machinery (i.e. SPIRE1/2 and FMN1), at the melanosome membrane. Here we suggest a model by which this might occur (Figure 7).

**Figure 7.**
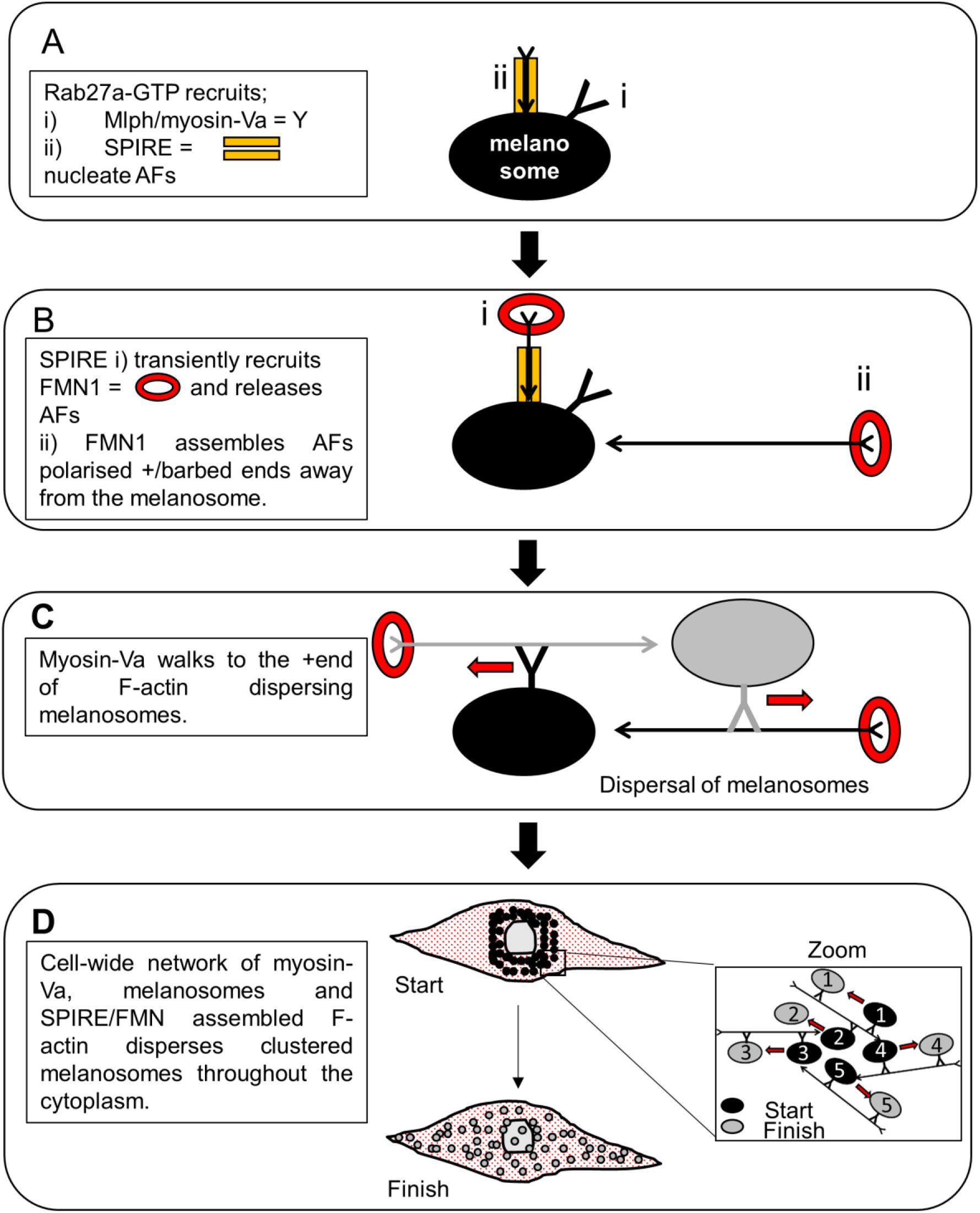
A model indicating how Rab27a could regulate AF driven melanosome transport by integrating the activity of the motor protein myosin-Va and track assembly proteins SPIRE1/2 and FMN1. See main text for details. In C) grey and black colours indicate AF and myosin-Va associated with melanosomes of the same colour. In D) grey and black colours indicate the position of numbered melanosomes at the start and finish of an episode of dispersive transport.

Firstly, we propose that active Rab27a recruits 2 effector complexes to melanosomes; i) myosin-Va (via interaction with the RBD of Mlph) and ii) SPIRE1/2 (via interaction with the C-terminus SFH domains of SPIRE1/2) (Figure 7A). [The recently identified myosin-Va:SPIRE1/2 interaction might also stabilise SPIREs at the melanosome and/or integrate their activity with myosin-Va ^36^]. Melanosomal SPIRE1/2 may then i) assemble (and later release) mini-AFs, via WH2:G-actin interaction, and ii) transiently recruit FMN1 to the membrane via SPIRE1/2-KIND:FMN1-FSI interaction (Figure 7B) ^46^. Interestingly, a recent study found that intramolecular SPIRE2 KIND/FYVE and the trans-regulatory FMN2-FSI/SPIRE2-KIND interactions were competitive ^31^. Thus association of SPIRE1/2-SFH domains with Rab27a may enhance SPIRE1/2-KIND:FMN1-FSI interaction. We suggest that transient interaction with SPIRE1/2 allows FMN1 to associate with the +/barbed ends of AF nuclei at melanosomes and elongate them. Thereby generating an AF network that overall is polarised with +/barbed ends oriented away from the melanosome membrane and towards the cell periphery (Figure 7B). Myosin-Va (attached to adjacent melanosomes) may then ‘walk’ toward the +/barbed ends of these AFs thereby moving melanosomes away from one another (Figure 7C).

Extrapolating to the cellular level we propose that globally this pattern of myosin-Va/AF dependent inter-organelle repulsion could drive and sustain dispersal of melanosomes from one another, resulting in their dispersal throughout the cytoplasm as seen in wild-type melanocytes (Figure 7D). We suggest that such an inter-connected, three-dimensional network of organelles, tracks and motors could allow cell-wide coupling of local organelle associated force generators (i.e. myosin-Va motors and FMN1/SPIRE1/2 AF assembly machinery). This arrangement could allow the generation of sufficient force to rapidly disperse these large, rigid and numerous organelles, and maintain this distribution, more efficiently than conventional transport models, which envisage single organelles pulled along single tracks by motors independently of other cargo, motors and tracks. Consistent with this the sarcomere of muscle cells and stress fibres of migrating fibroblasts represent alternative arrangements of myosin and AFs that allow the amplification of force generated by myosin-II motors to drive cell shape changes and movement ^47,48^. Meanwhile recent work in oocytes reveal that myosin-Vb, SPIRE1/2 and FMN2 form an AF meshwork that drives rapid, long-range outward flow of Rab11 positive vesicles to the plasma membrane, independently of MTs ^22,49^. With our data this suggests that similar AF-dependent organelle transport systems might be widespread in somatic mammalian cells.

### Stable attachment of SPIRE, but not FMN1, to melanosomes enhances its transport function

To test the model, we investigated the prediction that SPIRE1/2 and FMN1 function proximal and distal to the melanosome membrane (Figure 7A-B). For this we probed the functional effect of tethering the active fragments of SPIRE2 (KW) and FMN1 (FH1-FH2) to melanosomes by expressing them as fusions to the melanosome-targeted, non-functional, Rab27a^SF1F4^ mutant (hereafter KW-Rab27a and FH1-FH2-Rab27a) in SPIRE1/2 depleted melan-a or melan-f cells ^50^. We found that KW-Rab27a localised to melanosomes and restored their dispersion in SPIRE1/2 depleted cells to a greater extent than SPIRE2-KW alone, indicating that association with melanosomes enhances its function (Figure 5A, B (high magnification inset), C; mean % cells with dispersed or hyper-dispersed melanosomes KW-Rab27a = 98.01 +/- 2 %). This supports the suggestion that SPIRE1/2 assemble AF nuclei in close proximity to the melanosome membrane. Interestingly these experiments also revealed that fusion of KW to Rab27a reduced its tendency to drive hyper-dispersion of melanosomes (Figure 5C). This further supports the idea that differences in interaction of SPIRE1/2 with Rab27a underlie differences in their ability to disperse melanosomes.

In contrast, although FH1-FH2-Rab27a localised to melanosomes in melan-f cells and increased melanosome dispersion compared with GFP, it was significantly less effective than intact FMN1 or the FH1-FH2-FSI fragment, indicating that stable association with melanosomes reduces FMN1 function (Figure 6A, B (high magnification inset), C; mean pigment area (% total), FH1-FH2-Rab27a = 55.42 +/- 17.49 %). With data on the importance of the FSI motif supports the suggestion that FMN1 only transiently associates with melanosome associated SPIRE1/2.

### Membrane targeting of FMN family formins reduces their function in melanosome transport

To further investigate the functional importance of the subcellular positioning of the SPIRE1/2 and FMN1 AF assembly machinery we tested the ability of FMN1 related protein, FMN2, to rescue melanosome transport in melan-f cells. Like FMN1, FMN2 contains FH1 and FH2 domains and the C-terminus FSI, but in contrast to FMN1, the N-terminus amino acid sequence of FMN2 contains a putative N-myristoyltransferase (NMT) recognition sequence (Figure 8A) ^51^. This suggests that FMN2 may be post-translationally myristoylated and targeted to the cell membranes like other myristoylated proteins e.g. c-Src ^52^. Thus, FMN2 may generate AF close to cellular membranes i.e. distant from the perinuclear clustered melanosomes in melan-f cells, rather than throughout the cytoplasm as appears to be the case for cytoplasmically located FMN1 (Figures 1F, 6B and 8B). We found that both full length FMN2 (hereafter GFP-FMN2) and the FMN2 FH1-FH2-FSI fragment expressed as fusions to the C-terminus of GFP were distributed throughout the cytoplasm and restored melanosome dispersion with similar efficiency to GFP-FMN1 (Figure 8; mean pigment area (% total); GFP-FMN1 = 84.11 +/- 9.544 %, GFP-FMN2 = 81.72 +/- 8.011 % and GFP-FMN2 FH1-FH2-FSI = 86.42 +/- 10.84 %). Meanwhile, although FMN2 fused to the N-terminus of GFP (FMN2-GFP) dispersed melanosomes to a greater extent than GFP alone, it did so with significantly lower efficiency than GFP-FMN2 (Figure 8; mean pigment area (% total); GFP = 44.17 +/- 9.31%, FMN2-GFP = 58.37 +/- 7.818%). We also observed that FMN2-GFP localised to discreet cytoplasmic structures to a greater extent than did GFP-FMN2, consistent with its potential post-translational myristoylation and membrane targeting (Figure 8B, see high magnification inset images).

**Figure 8.**
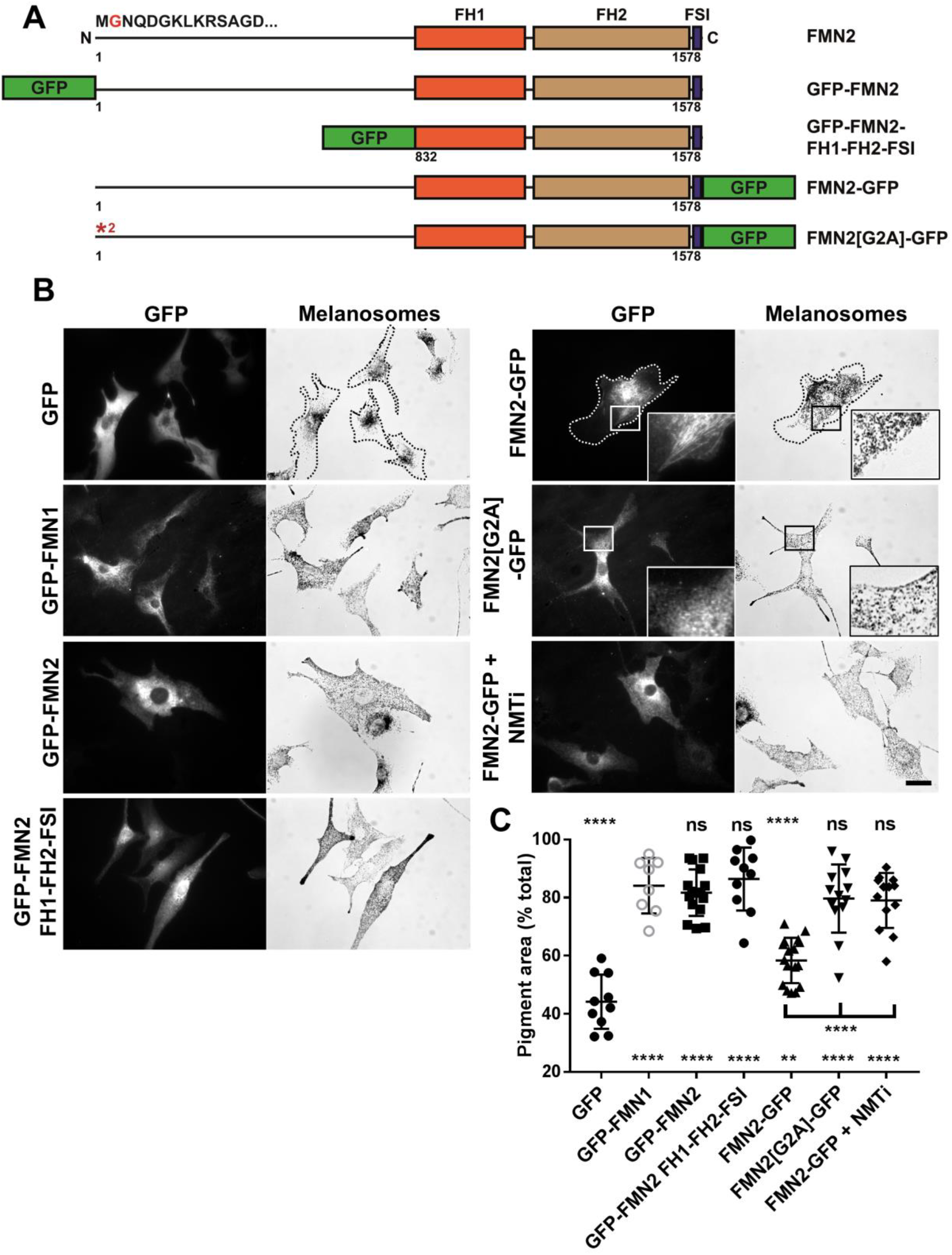
Perturbation of myristoylation of FMN2 enhances its ability to rescue melanosome transport in FMN1 deficient melanocytes. melan-f cells were infected with adenoviruses expressing the indicated proteins in the presence/absence of NMTi IMP-1088. Cells were fixed and stained for immunofluorescence 7 hours later using anti-GFP antibodies, and the distribution of expressed protein (GFP) and melanosomes was recorded using a fluorescence microscope (see Experimental procedures). A) A schematic representation of the domain structure of murine FMN2 and the composition of truncations and chimeric proteins used in functional studies (B-C). Red text indicates the site of the glycine [2] that is predicted to undergo myristoylation. Numbers indicate amino acid boundaries. B) Representative images of cells expressing each protein. Dotted lines in FMN2-GFP images highlight the borders of cells. Scale bar represents 20 μm. C) A scatter-plot showing the extent of pigment dispersion (pigment area (% total)) in populations of cells in each condition. n = 10 (GFP), 8 (GFP-FMN1), 16 (GFP-FMN2), 10 (GFP-FMN2-FH1-FH2-FSI) 17 (FMN2-GFP), 13 (FMN2[G2A]- GFP) and 13 (FMN2-GFP + NMTi (IMP-1088 100nM)). **** and ** indicate significant difference p = < 0.0001 and p = < 0.01 as determined by one-way ANOVA. n.s. indicates no significant difference. Significance indicators above and below the data indicate differences between each condition and the positive (GFP-FMN1) and negative (GFP alone) controls. Results shown are representative of 3 independent experiments. *FH*, formin homology; *FSI*, Formin-SPIRE interaction sequence.

To investigate whether myristoylation, and membrane targeting, could be the basis of the reduced function of FMN2-GFP versus GFP-FMN2 we tested the effect of blocking FMN2-GFP myristoylation by two strategies. Firstly, we tested the function of the FMN2-[G2A]-GFP mutant in which the glycine at position 2, predicted to be myristoylated, was replaced with non-myristoylatable alanine. Secondly, we tested the effect of a recently identified small molecule NMT inhibitor (IMP-1088) on the function of FMN2-GFP in melanosome transport (Figure 8) ^53^. We found that both strategies redistributed FMN2-GFP to a cytoplasmic distribution pattern that was similar to that seen for GFP-FMN2 and GFP-FMN1 and significantly enhanced its function in transport (Figure 8; mean pigment area (% total); FMN2-[G2A]-GFP = 79.71 +/- 11.72%, FMN2-GFP + NMTi = 79.03 +/- 9.488%). These observations suggest that myristoylation and tethering to membranes reduces the efficiency of FMN2 in melanosome transport. This is consistent with the prediction of the model which envisages that the less spatially restricted distribution of FMN1, vs SPIRE1/2, would allow it to transiently associate with SPIRE1/2 at the melanosome membrane and extend SPIRE1/2 assembled AF nuclei from the melanosome membrane (Figure 7).

### FMN1 assembles a melanosome associated AF network in melanocytes

As a final test of the hypothesis that SPIRE1/2 and FMN1 assemble AFs that are essential for transport we used field emission scanning electron microscopy (FESEM) to compare the amount and distribution of AFs associated with melanosomes in melan-a and melan-f cells. High magnification FESEM images revealed that dispersed melanosomes in melan-a cells were surrounded by a dense meshwork of AFs (Figure 9A, C, E). In striking contrast relatively few AFs were seen in the vicinity of melanosomes in melan-f cells (Figure 9B, D, F). Interestingly, close observation revealed a relative abundance of short AFs (or stumps) on melanosomes in melan-f, compared with melan-a, cells (Figure 9F arrows). These may represent SPIRE1/2 assembled AF nuclei that persist in these cells due to the lack of FMN1. These observations further support the hypothesis that SPIRE1/2 and FMN1 assemble AFs adjacent to melanosomes that may be used by myosin-Va to disperse these organelles.

**Figure 9.**
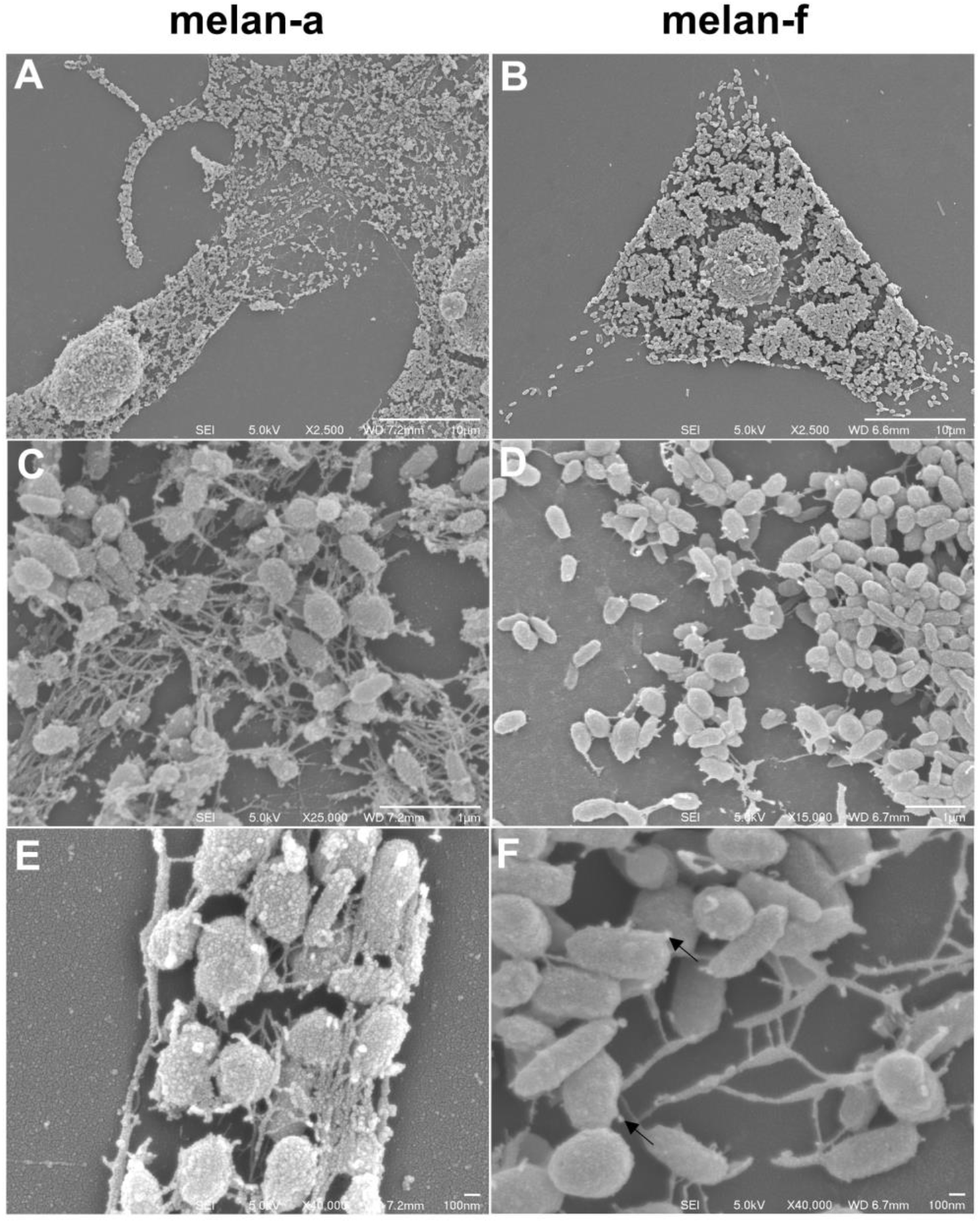
Scanning electron microscopy reveals differences in the abundance of melanosome associated AFs in wild-type and FMN1 deficient melanocytes. Wild-type (melan-a) and FMN1 deficient (melan-f) cells were prepared for scanning electron microscopy (SEM; see Experimental procedures). A), C) and E) are melan-a while B), D) and F) are melan-f cells. Arrows in F) indicate short actin filaments (or stumps) that are more abundant in melan-f compared with melan-a cells.

## Discussion

In this work we investigated the mechanism of myosin-Va-dependent dispersion of melanosomes in melanocytes, specifically focusing on the role of dynamic/latrunculin-A sensitive AFs that we previously identified as essential for this. We discovered that the FMN1 and SPIRE1/2 are the architects of this dynamic AF population. We also found that SPIRE1/2 are Rab27a effector proteins that may associate with melanosomes in a Rab27a dependent manner. Our work also highlighted functional divergence between SPIRE1/2 e.g. differences in affinity of Rab27a interaction, and consequent differences in cellular distribution and capacity to dispersed organelles. Overall our data suggest that SPIRE1 is the predominant AF nucleator in melanosome transport. Based on these findings we suggest that SPIRE1/2 and FMN1 assemble a dynamic AF network that may link melanosomes and be polarised such that the -/pointed ends are proximal to melanosomes, as a result of Rab27a dependent recruitment of SPIRE1/2 to melanosomes, while the +/barbed ends extend away from melanosomes into the cytoplasm. Thus, the barbed end directed motor myosin-Va can disperse melanosomes from one another by moving along these SPIRE1/2 and FMN1 assembled AFs.

Several pieces of data presented here, and previously, support this possibility. Firstly, melan-f cells show significantly fewer AFs in near melanosomes compared with melan-a cells (Figure 9). Secondly, fusion of active SPIRE1/2 and FMN1 to melanosomes via the Rab27a^SF1F4^ mutant boosts the function of active SPIRE2-KW and reduces the function of active FMN1-FH1-FH2, consistent with stable and transient modes of association of SPIRE1/2 and FMN1 with melanosomes. Thirdly, consistent with the predicted requirement for proximity between melanosomes and AFs, re-targeting of FMN (FMN2-GFP) to membranes via myristoylation reduces its function (Figure 8). Fourthly, we previously found that processive +/barbed end directed transport activity of myosin-Va was essential for its function in melanosome dispersion ^15^.

We suggest that our data have broader implication for transport in the many other cell types in which Rab27, related Rab proteins, SPIRE1/2, FMN and myosin-V proteins are expressed. In particular we note the similarity between the proteins working together in AF-dependent melanosome transport (SPIRE1/2, FMN1, Rab27a and myosin-Va) and those previously shown to regulate the assembly and dynamics of a vesicle associated AF meshwork that drives asymmetric spindle positioning during meiosis in mouse oocytes (SPIRE1/2, FMN2, Rab 11 and myosin-Vb) ^21,22^. The finding that strikingly similar groups of proteins drive strikingly similar activities i.e. rapid, long-range AF-dependent organelle transport, in different cell types suggests that related machineries may drive long-range organelles transport in a wide variety of other animal cell types. Consistent with this Rab27 is expressed wide variety of specialised ‘secretory’ cell type where it is associated with organelles whose contents are secreted into the extracellular space after their initial transport to the cortex from the cell body e.g. inflammatory mediator containing granules in mast cells, lytic granules in cytotoxic lymphocytes and exosome containing MVBs in cancer cells ^54-56^. The oligodactylism phenotype of the FMN1 knockout mouse strongly supports the possible function of FMN1 in secretory transport processes, since proper limb development requires a concerted communication between cells, which is vitally dependent on a circuit of secreted proteins ^37^. Also, the behavioural phenotypes found in FMN2 and SPIRE1 mutant mice may be the cause of altered secretion of neuropeptides or growth factors ^57,58^. Thus, it will be very interesting to test whether Rab27a contributes in a similar way to co-ordinating the AF-dependent transport of other organelles in other cell types.

### Experimental procedures

#### Derivation and maintenance of cultured cells

Cultures of immortal FMN1-deficient melanocytes (melan-f) were derived essentially as described previously ^59^. In brief mice carrying a previously generated FMN1 loss of function allele (FMN1^pro^) were crossed with Ink4a-Arf mutant mice in order to generate pups homozygous for the FMN1^pro^ mutant allele and heterozygous for Ink4a-Arf mutant allele. Genotyping of the embryos was as previously described ^37,59^. Melanocytes were then derived from the dorsal skin of mutant embryos as previously described ^60^. Cultures of immortal melan-f, melan-ash, melan-ln and melan-a melanocytes and HEK293a were maintained, infected with adenovirus expression vectors and transfected with siRNA oligonucleotides as described previously 42,61. The melanocyte cell lines described here are available from the Wellcome Trust Functional Genomics Cell Bank http://www.sgul.ac.uk/depts/anatomy/pages/WTFGCB.htm.

#### Transfection of cultured cells

Transfection of cultured cells with siRNA and plasmid DNA was as previously described ^61^. NT, Rab27a specific siRNA were as previously described ^62^. Sequence of siRNA for SPIRE1/2 and FMN1 are indicated in Table 1. siRNA oligonucleotides were purchased from Dharmacon Thermo-Fisher and Sigma Genosys UK.

**Table 1.**
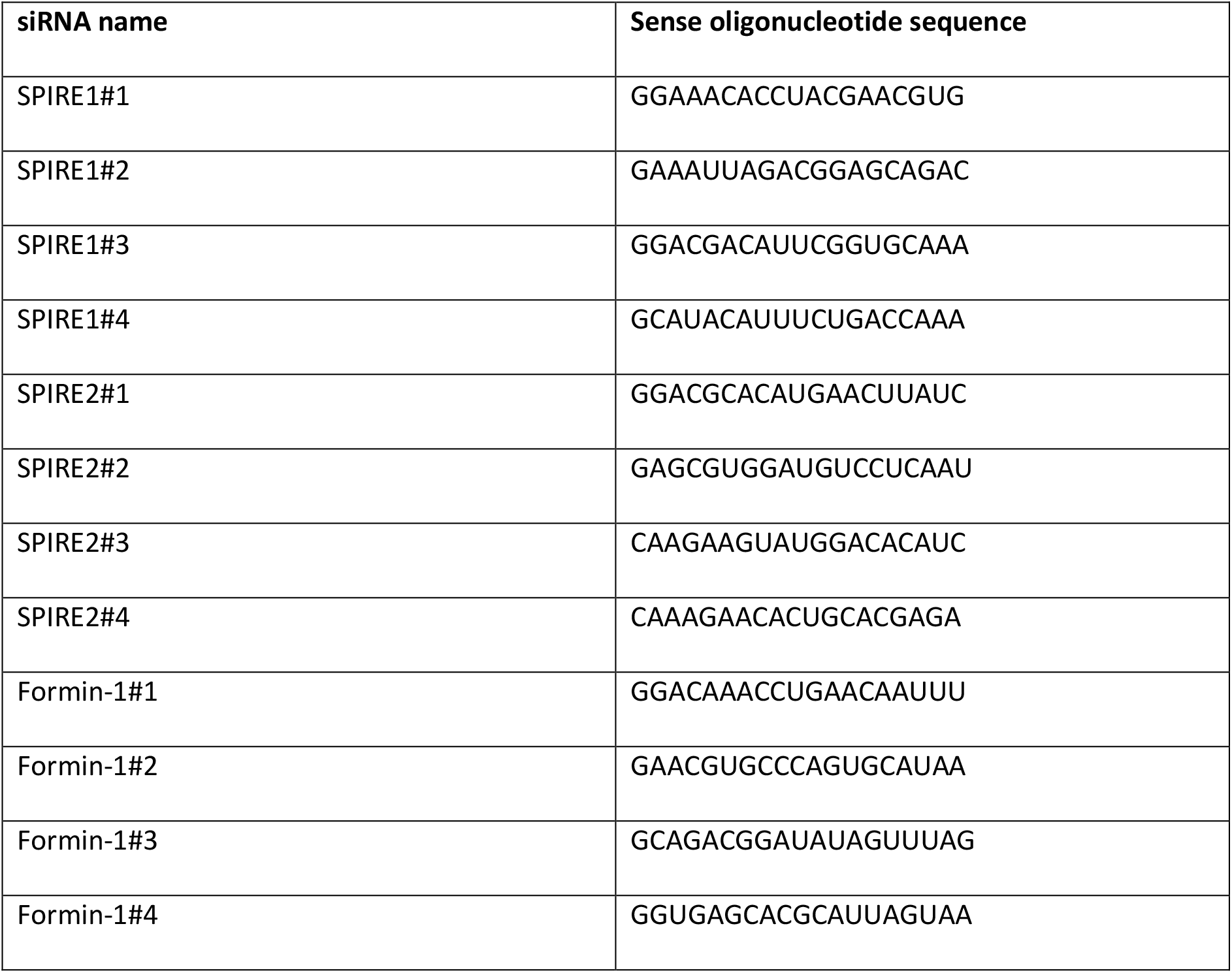
siRNA oligonucleotides used in this study.

#### Quantitative real-time PCR

Primers and the probes for Q-RT-PCR targets (from Sigma Genosys, Cambridge, UK) were designed using Primer Express software (Life Technologies). Probes were labelled at the 5’- and 3’- ends with fluorophore 6-FAM (6-carboxyfluorescein) and quencher TAMRA (tetramethylrhodamine), respectively (Table 2). To generate mRNA samples pools of melan-a cells grown in 6-well plates (1 × 10^5^/well) were transfected with siRNA in triplicate as described previously 61. 72 hours later cells were harvested and mRNA extracted using the RNeasy Mini RNA extraction kit (Qiagen). cDNA was generated using Moloney Murine Leukemia Virus M-MLV reverse transcriptase (Promega) using random primers. To generate a standard curve of signal:template concentration for each Q-RT-PCR assay a pool containing 5 % of each of the cDNA samples analysed was generated.

**Table 2.**
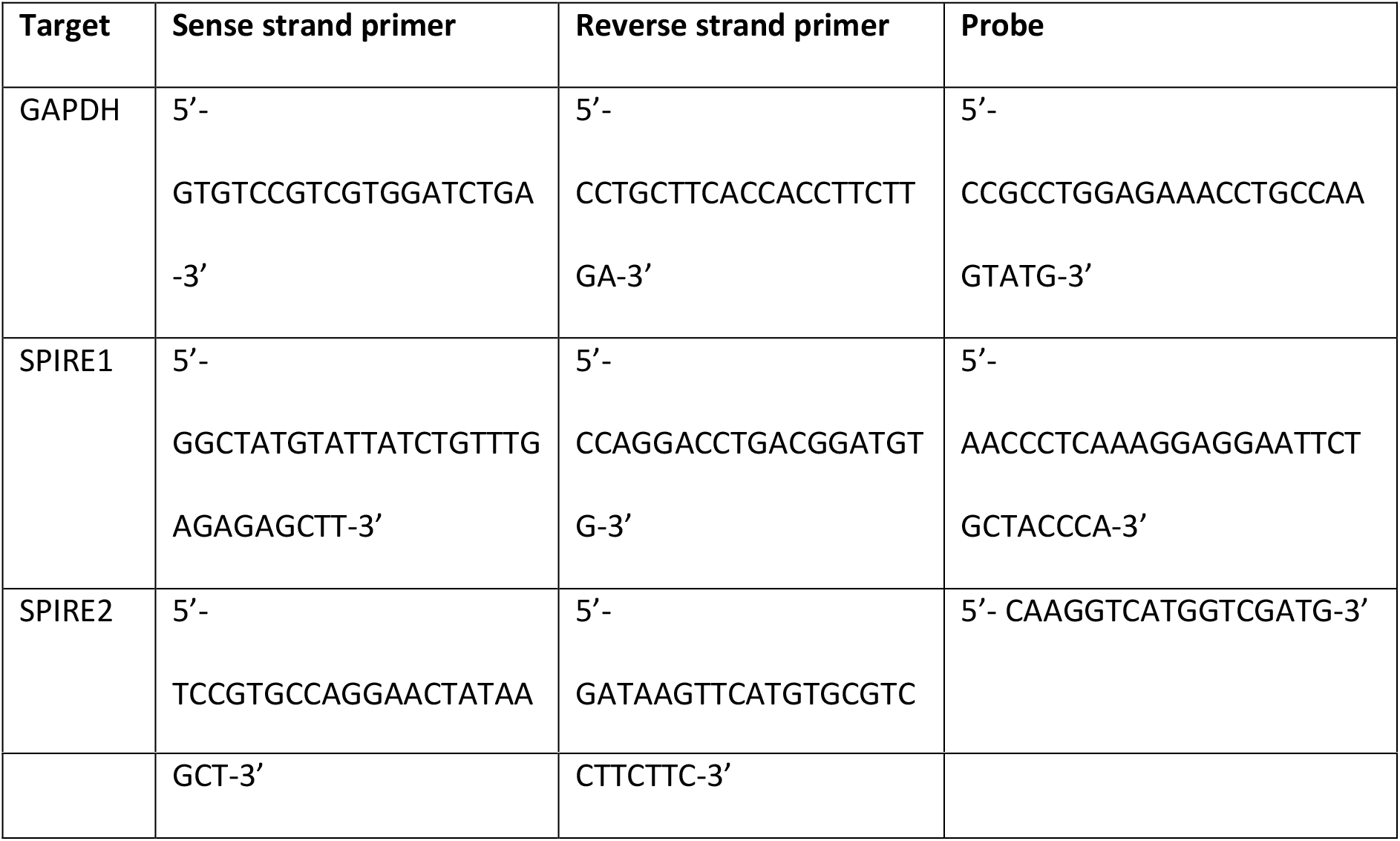
Taqman primers and probes used in this study.

This pool was serially diluted in DEPC water (1:4, 1:16, 1:64, 1:256) and these were used as template in target gene and GAPDH Q-RT-PCR assays. [The shape of the standard curve indicates the relationship between signal and template concentration. For all assays standard curves gave straight lines with R^2^ > 0.99 indicating that there is a linear relationship between signal and template]. To measure the expression of targets in siRNA transfected cells each neat cDNA was diluted 1:32 in DEPC water and the following reagents were added per well of a 96-well plate: 6.5 μl TaqMan Fast 2X PCR Master Mix (Life Technologies); 0.4 μl forward primer (10 μM); 0.4 μl reverse primer (10 μM); 0.25 μl probe (10 μM); 3 μl cDNA; 2.45 μl DEPC water. For each sample, 3 technical repeats were performed. Reaction plates were sealed with optically clear adhesive film, centrifuged, and qRT-PCR performed using a StepOnePlus Real-Time PCR system (Applied Biosystems) using the ‘fast’ mode. CT values for each reaction were determined by the StepOne software. The slope (S), intercept (I) and R^2^ values were calculated for the standard curve of each Q-RT-PCR assay. CT values from siRNA transfected samples were then processed to generate a ‘quantity value’ for each CT value as follows; 1) (CT-I)/S=LQ, 2) 10LQ=Q, 3) Qx(1/MNT)= GOIP (where MNT= mean non-targeted quantity value) and 4) GOIP/GP = normalised expression of target relative to GAPDH (where GP is the normalised quantity value for the GAPDH primer). Expression of FMN1 was detected using Taqman assay Mm01327668_m1 from Thermo-Fisher Scientific, UK.

For measurement of the expression levels of SPIRE and FMN genes the RNA isolation kit of Macherey-Nagel was used. To reduce amount of melanin and increase purity of total RNA step two of the protocol was repeated. In order to quantify the amount of total RNA isolated, spectrophotometric determination was used. cDNA was generated employing the Qiagen QuantiNova^®^ Reverse Transcription Kit (according the manufacturer’s protocol). An internal control RNA was used to verify successful reverse transcription. PCR primer sets were designed using primer3, Primer-Blast and were constructed by Sigma-Aldrich. The primer sets and templates used are shown in Table 3. Generated cDNA from melan-a total RNA and plasmid DNAs as positive controls were used as templates. For standard PCRs Q5^®^ High-Fidelity DNA Polymerase from New England Biolabs and the corresponding protocol were used. For absolute quantification an external plasmid-cDNA standard was used to create a standard curve with known copy number concentrations. Linearised plasmid DNA was serially diluted in water. The copy number of standard DNA molecules could be determined by the following formula: ((X g/μl DNA)/[plasmid length in basepairs × 660])× 6.022 × 10^23^ = Y molecules/μl. The absolute quantification by qPCR was performed with the Rotor-Gene Q (Qiagen) thermocycler. For each reaction triplets were performed, and CT values were determined with the help of the Qiagen Rotor-Gene Q Series Software. CT values of the samples are compared to the standard curve with known concentrations to calculate the amount of target in the samples. For this purpose, the QuantiNova^®^ SYBR^®^ Green PCR Kit and the associated protocol were employed.

**Table 3.**
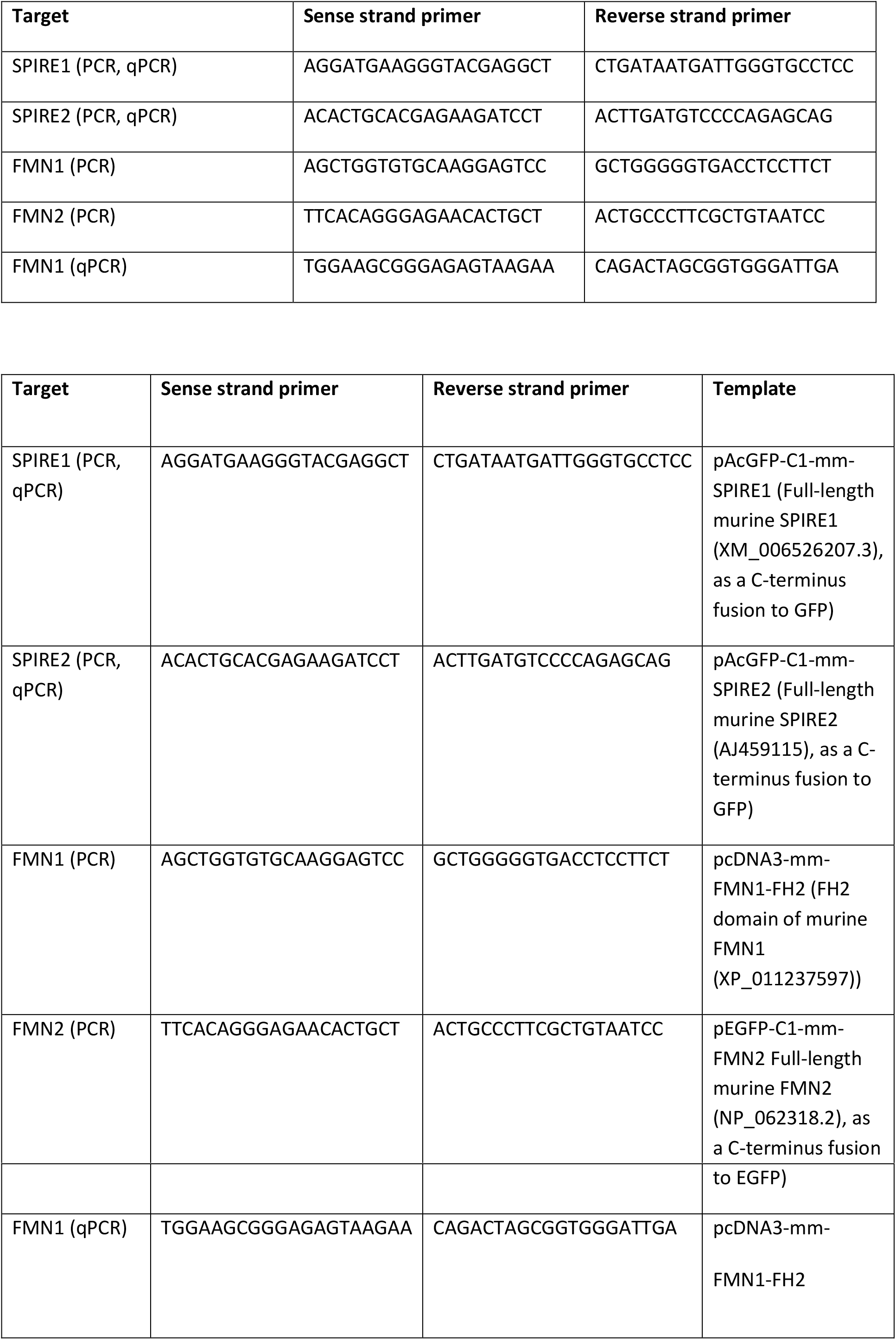
Primers and control templates used in qRT-PCR experiments.

**Table 4.** Prokaryotic and eukaryotic expression vectors used in this study. Unless otherwise stated vectors allow expression of protein in mammalian cells and were generated in this study. pENTR vectors were used to generate adenoviruses as previously described ^42^.

| Vector | Expressed protein |
| --- | --- |
| pENTR-EGFP | EGFP alone <sup>42</sup>. |
| pENTR-EGFP-C2-mm-FMN1, FH1-FH2-FSI, N-term, ΔFSI, FH2-FSI | Murine full-length FMN1 and aa849-1430, aa1-859, aa1-1394, aa985-1430 fragments as a fusion to the C-terminus of EGFP. Generated by PCR and sub-cloning from pEGFPC2-FMNiso1b (Addgene #19320) (XP_011237597) <sup>69</sup>. |
| pENTR-EGFPC1-Myc-hs-SPIRE1 | Human SPIRE1 (KIAA1135) as a fusion to the C-terminus of EGFP-Myc-epitope. |
| pENTR-EGFPC1-Myc-hs-SPIRE2 | Human SPIRE2 (AJ422077) as a fusion to the C-terminus of EGFP-Myc-epitope. |
| pGEX4T1-rn-Rab27a-Q78L and -T23N | Active and inactive mutant forms of rat Rab27a (P23640) in *E.coli* as fusions to the C-terminus of GST. |
| pcDNA3-Myc-hs-SPIRE1, KWM and MSFH | Full-length human SPIRE1 (KIAA1135), aa2-524 and aa388-742 fragments as a fusion to the C-terminus of the Myc-epitope. |
| pcDNA3-Myc-hs-SPIRE2 and MSFH | Full-length human SPIRE2 (AJ422077) and aa361-714 fragment as a fusion to the C-terminus of the Myc-epitope. |
| pAcGFP1-C1-hs-SPIRE1-MSFH, SFH, SF, FYVE and SB | Full-length human SPIRE1 (KIAA1135), aa388-742, aa525-742, aa525-697, aa573-697 and aa525-572 fragments as a fusion to the C-terminus of AcGFP1. |
| pCMVMYC-mCherry | Myc-mCherry fusion protein <sup>70</sup>. |
| pENTR-V5-vYn Myc-hs-SPIRE1 and hs-SPIRE2 | Full-length human SPIRE1 and SPIRE2 as a fusion to the C-terminus of the V5 epitope tag and an N-terminus fragment (ACE74713) of Venus YFP. |
| pENTR-V5-vYc- rn-Rab27a, T23N, Q78L, N133I | Rat Rab27a as a fusion to the C-terminus of the V5 epitope tag and an C-terminus fragment (AM779184.1) of Venus YFP. |
| pAcGFP1-C1-mm-Mlph RBD | Rab27a binding domain of murine Mlph (aa 1-150; NP_443748.2) as a C-terminus fusion to GFP. |
| pENTR-EGFPC2-mm-MyoVa1-920aa-SytI2-a RBD | Active S1 (motor-lever) fragment of murine myosin-Va fused at the N-terminus to GFP and the C-terminus to the Rab27a binding domain of murine SytI2-a, described previously <sup>15</sup>. |
| pENTR-EGFPC2-mm-MyoVa1-920aa-SPIRE1/2 | As above but with SytI2-a replaced with either murine SPIRE1 or SPIRE2. Murine SPIRE coding sequence was PCR amplified from EST clones IRAVp968A10102D (Q52KF3.1) and IRAVp968C09162D (NP_758491.1) obtained from Source Bioscience, Nottingham, UK. |
| pENTR-EGFPC1-hs-SPIRE2 MSFH, KW, WMSFH | Human SPIRE2 (AJ422077) protein fragments; aa |
| and KMSFH | 361-714, aa1-370, aa230-714, aa2-230 with 361-714. |
| pENTR-EGFPC1 hs-SPIRE2-KW-Rab27a | Human SPIRE2 (AJ422077) aa1-370 fused to the C-terminus of EGFP and the N-terminus of Rab27aSF1F4<sup>50</sup>. |
| pENTR-EGFPC2-mm-FMN1-FH1-FH2-Rab27a-sf1f4#4 | Murine FMN1 fragment aa840-1401 fused to the C-terminus of EGFP and the N-terminus of Rab27aSF1F4<sup>50</sup>. |
| pENTR-EGFPN3-mm-FMN2, pENTR-EGFPC1-mm-FMN2 | Murine FMN2 full length (NP_062318.2) fused to the N- and C-termini of EGFP. |
| pENTR-EGFPC1-mm-FMN2-FH1-FH2-FSI | Murine FMN2 aa854-1587 fused to the C-terminus of EGFP. |
| pEGFPC3-rn-Rab27a | Rat Rab27a as a fusion to the C-terminus of EGFP<sup>71</sup>. |
| pmRuby3-C1-rn-Rab27a | Rat Rab27a as a fusion to the C-terminus of mRuby3. |
| pEGFP-C2-mm-MyoVa-CC-GTD | Murine myosin Va fragment (aa 1260-1880) as a fusion to the C-terminus of EGFP<sup>72</sup>. |

#### Immunoblotting

Immunoblotting was performed as described previously ^61^ (Pylypenko et al., 2016) using rabbit anti-FMN1 (diluted 1:1000), goat anti-GAPDH (Sicgen Ab0049-200; 1:5000), anti-GFP (Living Colors A.v. peptide antibody, rabbit polyclonal, 1 mg/ml; Clontech) and anti-c-Myc (9E10, mouse monoclonal, 0.4 mg/ml; Santa Cruz Biotechnology) primary antibodies, and IRDye 800CW conjugated secondary antibodies (Odyssey 926-32214; 1:10000) or horseradish peroxidase linked anti-rabbit IgG (from donkey) and anti-mouse IgG (from sheep) secondary antibodies (1:5000, GE Healthcare Life Sciences, Freiburg, Germany). Signal was detected using a Li-Cor infrared scanner (Odyssey) or by chemiluminescence (Luminata Forte Western HRP substrate; Merck Millipore, Darmstadt, Germany) recorded with an Image Quant LAS4000 system (GE Healthcare Life Sciences).

#### *Multiple protein sequence alignments and phylogenetic tree* generation

Multiple sequence alignments for C-terminal regions of chosen SPIRE1 and SPIRE2 and N-terminal Mlph and Myrip amino acid sequences, respectively, were performed using the Clustal Omega Multiple Sequence Alignment tool (EMBL-EBI, Hinxton, UK) and subsequent manual confinements. Respective cDNA sequences were obtained from the NCBI Gene database with the following accession numbers: Homo sapiens (Hs)-SPIRE1: NM_020148.2, Hs-SPIRE2: AJ422077, Mus musculus (Mm)-SPIRE1: XM_006526207.3, Mm-SPIRE2: AJ459115, Gallus gallus (Gg)-SPIRE1: XM_419119.4, Gg-SPIRE2: XM_004944321.1, Hs-Mlph: NM_024101.6, Mm-Mlph: NM_053015.3, Gg-Mlph: NM_001115080.1, Hs-Myrip: NM_001284423.1, Mm-Myrip: NM_144557.5, Gg-Myrip: XM_015281725.1.

Generation of phylogenetic trees for Rab27 interacting proteins was based on multiple amino acid sequence alignments using Clustal Omega as described above and manual confinements. Respective cDNA sequences were obtained from the NCBI Gene database with the following accession numbers: Hs-RIMS1: NM_014989.5, Hs-RIMS2: NM_001100117.2, Hs-RPH3A (Rabphilin-3A): NM_001143854.1, Hs-RPH3AL (Noc2): NM_006987.3, Hs-Mlph (Slac2-a): NM_024101.6, Hs-EXPH5 (Slac2-b): NM_015065.2, Hs-Myrip (Slac2-c): NM_001284423.1, Hs-SYTL1 (Slp1): XM_005246022.3, Hs-SYTL2 (Slp2-a): NM_032943.4, Hs-SYTL3 (Slp3-a): NM_001242384.1, Hs-SYTL4 (Slp4, granuphilin): NM_080737.2, Hs-SYTL5 (Slp5): NM_001163334.1, Hs-SPIRE1: NM_020148.2, Hs-SPIRE2: AJ422077. Evolutionary analyses and phylogenetic tree formation were conducted in MEGA7 ^63^. The evolutionary history was inferred by using the Maximum Likelihood method based on the JTT matrix-based model ^64^. The bootstrap consensus tree inferred from 500 replicates is taken to represent the evolutionary history of the taxa analysed ^65^. Branches corresponding to partitions reproduced in less than 50 % bootstrap replicates are collapsed. The percentage of replicate trees in which the associated taxa clustered together in the bootstrap test (500 replicates) are shown next to the branches ^65^. Initial tree(s) for the heuristic search were obtained automatically by applying Neighbor-Join and BioNJ algorithms to a matrix of pairwise distances estimated using a JTT model, and then selecting the topology with superior log likelihood value. The analysis involved 14 amino acid sequences. There was a total of 188 positions in the final dataset.

#### Cloning of bacterial and mammalian protein expression vectors

Expression vectors were generated by standard cloning techniques using Accuprime Pfx (Thermo Fisher Scientific) or Pfu (Promega, Mannheim, Germany) DNA polymerases, restriction endonucleases and T4 DNA ligase (both New England Biolabs, Frankfurt am Main, Germany). Point mutants were generated using the In-Fusion HD cloning kit (TakaraBio/Clontech). DNA sequencing was carried out by LGC Genomics (Berlin, Germany) and Source Bioscience (Nottingham, UK). Table 3 shows details of vectors used in this study. Amino acid boundaries are related to the following cDNA sequences: human SPIRE1 (NM_020148.2), human SPIRE2 (AJ422077), murine FMN1 (XP_011237597), murine FMN2 (NP_062318.2), murine myosin Va (XM_006510827.3) and murine Mlph (NP_443748.2).

#### Recombinant protein expression and purification

Recombinant GST-Rab27a-Q78L and GST-Rab27a-T23N proteins were expressed in *Escherichia coli* Rosetta bacterial cells (Merck Millipore, Novagen). Bacteria were cultured in LB medium (100 mg/l ampicillin, 34 mg/l chloramphenicol) at 37°C until an A600 nm of OD 0.6–0.8. Protein expression was induced by 0.2 mM Isopropyl-β-D-thiogalactopyranoside (IPTG; Sigma-Aldrich, Taufkirchen, Germany) and continued at 16–20°C for 18–20 hr. Bacteria were harvested and lysed by ultra-sonication. Soluble proteins were purified by an ÄKTApurifier system (GE Healthcare Life Sciences, Freiburg, Germany) using GSH-Sepharose 4B (GE Healthcare Life Sciences) for GST-tagged proteins and Ni-NTA beads (Qiagen, Hilden, Germany) for His6-tagged proteins, respectively, and size exclusion chromatography (High Load 16/60 Superdex 200; GE Healthcare Life Sciences). Proteins were concentrated by ultrafiltration using Amicon Ultra centrifugal filters (Merck Millipore) with respective molecular weight cut offs. The final protein purity was estimated by SDS-PAGE and Coomassie staining.

#### GST pull-down from HEK293 cell lysates

Hek293 cells were cultured and transfected with plasmid DNA as described previously ^36^. To ensure equal input of GFP-tagged or Myc-epitope-tagged SPIRE1 and SPIRE2 deletion mutants for GST pull-downs all SPIRE mutants were first expressed in HEK293 cells and respective protein expression levels were analysed by Western blotting (anti-GFP, anti-c-Myc). Protein bands were quantified using ImageQuant TL (GE Healthcare Life Sciences) and normalized to the lowest signal band. According to quantification the amount of cell lysates employed in subsequent pull-downs was adjusted. For pull-downs HEK293 cells were transfected with expression vectors encoding GFP-tagged and Myc-epitope-tagged SPIRE1 and SPIRE2 proteins. 48 hr post transfection, cells were lysed in lysis buffer (25 mM Tris-HCl pH 7.4, 150 mM NaCl, 5 mM MgCl2, 10 % (v/v) glycerol, 0.1 % (v/v) Nonidet P-40, 1 mM PMSF, protease inhibitor cocktail) and centrifuged at 20,000 × *g*, 4°C, 20 min to remove insoluble debris. For GST pull-down assays 65 μg GST-Rab27a-Q78L and GST-Rab27a-T23N, respectively, (25 μg GST control) was coupled to GSH-Sepharose 4B beads (1:1 suspension) for 1 hr, 4°C on a rotating wheel. Beads were washed twice with pull-down buffer (25 mM Tris-HCl pH 7.4, 150 mM NaCl, 5 mM MgCl2, 10% (v/v) glycerol, 0.1 % (v/v) Nonidet P-40) and subsequently incubated with cell lysates for 2 hr at 4°C on a rotating wheel. Here, 1 ml lysates with the lowest expressed protein was employed and lysates with higher protein abundance were diluted with pull-down buffer to 1 ml, according to prior quantification. Beads were washed four times with pulldown buffer and bound proteins were eluted with 1x Laemmli buffer, denatured at 95°C for 10 min and subsequently analysed by immunoblotting.

#### Quantitative GST pull-down assays

GST pull-down assays were performed as described above from HEK293 cell lysates with increasing concentrations of GST-Rab27a-Q78L protein. According to prior quantification the concentration of ectopic expressed GFP-SPIRE1-MSFH was approximately 50 nM. Cell lysates were pooled and equally distributed to beads coupled GST-Rab27a-Q78L protein. Beads were pelleted and the cell lysate supernatant was diluted 1:4 with aqua. dest. Each sample was allowed to adapt to 20°C for 10 min in a water bath. The concentration-dependent binding of Rab27a-Q78L to C-terminal GFP-SPIRE1-MSFH was determined by fluorospectrometric analysis using FluoroMax- 4 Spectrofluorometer (Horiba Jobin Yvon, Bensheim, Germany). The same experiment was performed employing GFP-Mlph-RBD expressed in HEK293 cells and purified GST-Rab27a-Q78L proteins. The AcGFP1 green fluorescent protein was excited at 488 nm (slit = 5 nm) and the emission at 507 nm was recorded (emission maximum, slit = 5 nm) with an integration time of 0.1 sec. The data were calculated as ’fraction bound’ (*y*) compared to the initial fluorescence signal without GST-Rab27a-Q78L protein coupled to beads.

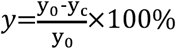

With I is fluorescence signal without GST-Rab27a-Q78L and *yC* is signal at corresponding GST-Rab27a-Q78L concentration. Furthermore, data were analyzed in SigmaPlot 12.3 software (Systat Software, Erkrath, Germany). Equilibrium binding data were fitted according to the equation

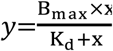

with *B_max_* representing the maximal amplitude, *K_d_* representing the equilibrium constant and *x* representing the concentration of GST-Rab27a-Q78L.

#### Microscopy and image analysis

Cells (1 × 10^4^) for immunofluorescence were cultured on 13 mm diameter 1.5 thickness glass coverslips (SLS, Nottingham, UK. 4616-324139) for at least 24 hours, prior to infection with adenovirus or transfection with plasmid DNA. Twenty-four hours later cells were paraformaldehyde fixed, stained and fluorescence and transmitted light images were then collected using a Zeiss LSM710 confocal microscope fitted with a 63x 1.4NA oil immersion Apochromat lens or a Zeiss Axiovert 100S inverted microscope fitted with 10x and 40x objective lenses and an Axiocam MR3 CCD camera. Antibodies and stains were used as indicated; mouse monoclonal anti-GFP (Roche 11814460001; 1:300) and goat anti-mouse Alexa568 labelled secondary antibodies (Invitrogen A-11011; 1:500), texas-red-phalloidin (Sigma P1951; 100 nM); anti-c-Myc antibody (9E10, mouse monoclonal, 2 mg/ml, Santa Cruz Biotechnology) and anti-TRP1 antibody (Ta99 mouse monoclonal, Santa Cruz Biotechnology sc-58438, 1:1000) and conjugated anti-mouse secondary antibodies (Cy5; from donkey, 3.25 mg/ ml, Dianova, Hamburg, Germany).

#### Analysis of melanosome dispersion/transport

For analysis of the effects of siRNA on melanosome distribution transfections were carried out in triplicate i.e. 3 wells of a 24-well plate for each siRNA in each experiment. 72 hours later phase contrast images of 3 different randomly-selected low power (using 10x objective) fields of cells in each well were captured using Axiovision 4.8 software associated with a Zeiss Axiovert 100S inverted microscope. Images were then randomised and the number of cells with clustered melanosomes in each field was counted by a researcher blinded to the identity of the siRNA transfected into each field of cells. Cells in which pigmented melanosomes were contained within the perinuclear cytoplasm (< 50 % of the total cytoplasmic area) were defined as containing clustered melanosomes. Measurement of the function of adenovirus expressed FMN1 and SPIRE1/2 proteins in melanosome transport was based on manual measurement of the proportion of cell area occupied by pigmented melanosomes as previously described ^42^.

#### Bimolecular fluorescence complementation

For bimolecular fluorescence complementation (BiFC) HEK293a cells were transfected with plasmids allowing expression of mCherry (control for transfection efficiency), vYNE-SPIRE1/2 (human SPIRE1/2 fused to the C-terminus of enhanced Venus YFP (accession CAO91538.1) N-terminus fragment (aa 1-173)) and vYCE-Rab27a wild-type and mutants (Rab27a (rat) fused to the C-terminus of enhanced Venus YFP C-terminus fragment (aa 155-239) using Fugene 6 transfection reagent (Promega E2691). Forty-eight hours later mCherry and YFP (BiFC) fluorescence was recorded from living cells in low power fields of view for each condition using the Zeiss Axiovert 100S microscope using the same exposure setting for each channel and each condition. Normalised BiFC signal was determined using ImageJ software. Briefly mCherry images were converted to binary and ROIs corresponding to transfected cells were saved to the regions manager. Mean vYFP for each ROI was extracted and normalised BiFC determined for each ROI by dividing the vYFP signal by the corresponding mCherry signal to normalise for differences in the level of transfection in each cell. Median BiFC signal for ROI was determine using Graphpad Prism 7 software (Graphpad, La Jolla, USA).

#### Colocalization analysis

Fixed cells were analyzed with a Leica AF6000LX fluorescence microscope, equipped with a Leica HCX PL APO 63x/1.3 GLYC objective and a Leica DFC350 FX digital camera (1392 × 1040 pixels, 6.45 μm × 6.45 μm pixel size) (all from Leica, Wetzlar, Germany). 3D stacks were recorded and processed with the Leica deconvolution software module. Images were recorded using the Leica LASX software and further processed with Adobe Photoshop and subsequently assembled with Adobe Illustrator. The extent of colocalization of myosin Va, Rab27a and SPIRE1 proteins at intracellular melanocyte membranes as well as the localization of Rab27a and SPIRE1 C-terminal fragments, respectively, at melanosome membranes was analyzed using the ImageJ (V2.0.0) plug-in Coloc2. Here, the colocalization rate is indicated by the Pearson’s Correlation Coefficient (PCC) as a statistical measure to unravel a linear correlation between the intensity of different fluorescent signals. A PCC value of 1 indicates a perfect colocalization, 0 indicates a random colocalization and a PCC value of −1 indicates a mutually exclusive localization of the analyzed signals. To take the noise of each image into account and to gain an objective evaluation of PCC significance, a Costes significance test was performed. To do so, the pixels in one image were scrambled randomly and the correlation with the other (unscrambled) image was measured. Significance regarding correlation was observed when at least 95 % of randomized images show a PCC less than that of the original image, meaning that the probability for the measured correlation of two colours is significantly greater than the correlation of random overlap ^66,67^.

#### Cytoskeleton preparations for FESEM

melan-a and melan-f cells were plated on 7 mm glass coverslips and grown for 24 hours. The medium was aspirated and the cells were extracted for 1 min with 0.25% Triton X-100 (Sigma, UK) and 5 μM phallacidin (Sigma) in cytoskeletal stabilization buffer (CSB: 5 mM KCl, 137 mM NaCl, 4 mM NaHCO3, 0.4 mM KH2PO4, 1.1 mM Na2HPO4, 2 mM MgCl2, 5 mM Pipes, 2 mM EGTA, and 5.5 mM glucose, pH 6.1; ^68^). After a quick wash in CSB, the samples were fixed with 1% glutaraldehyde (Sigma) in CBS with 5 μM phallicidin for 15 min. The coverslips were transferred to HPLC-grade water and incubated with 2% osmium tetroxide in water for 1 hour. After 3 fifteen-minute washes in water, the coverslips were dehydrated through successive 15 min immersion in 30, 50, 70, 90 and 100% alcohol solutions, followed by two 20 min incubation in methanol. The coverslips were carefully picked from the methanol solution and gently immersed vertically in HMDS (Hexamethyldisilazane, Sigma) in glass bottles for 30 seconds twice and left to air-dry for at least an hour before mounting them on 10 mm diameter specimen stubs. The coverslips were then coated with silver paint and sputter coated with a 5-6 nm layer of platinum an Edwards S150B sputter coater. Samples were imaged in a JEOL JSM-6700F scanning electron microscope by secondary electron detection.

## Acknowledgments

We thank Dr Nick Holliday (University of Nottingham, UK) for vYFP BiFC vectors, Dr Markus Dettenhofer (CEITEC - Central European Institute of Technology, Brno, Czech republic), Dr Volney Sheen (Harvard Medical School, Boston, USA) and Dr Monserrat Bosch (Institut Pasteur, Paris, France) for FMN1pro mutant mice and FMN1-specific antibodies, Dr Andrew Bennett (University of Nottingham, UK) and his team for help with Q-RT-PCR, Tim Self (confocal microscopy), Chris Gell (Image analysis), Sue Cooper, Carol Sculthorpe (general) (all University of Nottingham, UK), Annette Samol-Wolf (expression vector cloning, University Hospital Regensburg, Germany) for technical assistance, and Clare Futter (UCL, Institute of Ophthalmology, UK) for advice on electron microscopy studies. This work was supported by a Medical Research Council New Investigator Award to ANH (grant reference G1100063), Wellcome Trust Project Grant Awards (grant reference 108429/Z/15/Z to EVS), a Biotechnology and Biological Sciences Research Council (grant reference BB/F016956/1) and University of Nottingham funded PhD studentship awarded to CLR. DAB was funded by Biochemical Society Vacation Studentship. LM is funded through the Spanish Ministry of Economy. Industry and Competitiveness (MINECO) [BIO2015-70978-R]. EK and TW are funded by DFG SPP1464, KE 447/10-2” and DFG KE 447/18-1. FS is funded by DFG Research Training Group, GRK 2174 Award. NA is supported by a PhD studentship from Taibah University, Medina, Kingdom of Saudi Arabia.

## Author contributions

NA, CLR, TW, ELP, DAB, FS, AS, JR, MB, EK and ANH carried out experiments and analysed data. ANH wrote the manuscript. MB, ET and EK contributed to revision of the manuscript. MC, LM, PSG and EVS generated the immortal Fmn-1 deficient melanocyte cell line melan-f.

